# Applications of spatial models to ordinal data

**DOI:** 10.1101/2020.09.21.306001

**Authors:** Zhanyou Xu, Steven B. Cannon, William D. Beavis

## Abstract

Models have been developed to account for heterogeneous spatial variation in field trials. These spatial models have been shown to successfully increase the quality of phenotypic data resulting in improved effectiveness of selection by plant breeders. The models were developed for continuous data types such as grain yield and plant height, but data for most traits, such as in iron deficiency chlorosis (IDC), are recorded on ordinal scales. Is it reasonable to make spatial adjustments to ordinal data by simply applying methods developed for continuous data? The objective of the research described herein is to evaluate methods for spatial adjustment on ordinal data, using soybean IDC as an example. Spatial adjustment models are classified into three different groups: group I, moving average grid adjustment; group II, geospatial autoregressive regression (SAR) models; and group III, tensor product penalized P-splines. Comparisons of eight models sampled from these three classes demonstrate that spatial adjustments depend on severity of field heterogeneity, the irregularity of the spatial patterns, and the model used. SAR models generally produce better performance metrics than other classes of models. However, none of the eight evaluated models fully removed spatial patterns indicating that there is a need to either adjust existing models or develop novel models for spatial adjustments of ordinal data collected in fields exhibiting discontinuous transitions between heterogeneous patches.

## Introduction

Iron deficiency chlorosis (IDC) in soybeans is caused by the inability of the plant to utilize iron in the soil. Without enough iron, chlorophyll production is hampered, and the plant will suffer and possibly die. Symptoms of IDC are expressed in new leaf tissues of younger leaves, between the first and third trifoliate growth stages, vegetative stages V1 to V3 [1]. The typical symptoms of IDC is the yellowing of leaves, with interveinal chlorosis, while the veins remain green [2].

Soybeans are the second-most-planted field crop in the United States after corn, with a record-high of 90.16 million acres planted in 2016 [3]. It is estimated that IDC reduces yields in farmers fields by 20% each unit for each level of increased chlorosis scores. IDC is scored by field breeders on a 1 (no yellowing symptoms) to 5 (severe yellowing of leaves, and the plant dies) scale [4]. Soybean planting acreage in IDC-prone regions has increased from 1979 to 2017, with a 160% increase of soybean production areas into IDC-prone regions in the past 30 years [5]. This increase of soybean production area into IDC-prone regions has led to yield losses of 340 million tons, worth an estimated $120 million per year [6].

In the primary soybean production regions, soil micro-environmental variation results in location heterogeneity for iron deficiency. Thus, it is difficult to find large and uniform fields of calcareous soil that can be used to evaluate IDC in typical field plots, resulting in more experimental error than is desirable for selecting resistant genotypes [7]. Environmental conditions for soybean to develop IDC symptoms are ephemeral, usually existing for a couple weeks during the V1 to V3 stages of soybean development. Fields chosen for IDC testing are selected based on both historical IDC pressure records and potential for IDC conditions detected at the time of planting. In fields known to exhibit IDC, the exact locations within the fields may change from year to year, depending on rainfall prior to planting, and the rate at which soil moisture evaporates in the early growing season. Within a testing site, IDC pressure usually varies either by ranges and rows from year to year, leading to different levels and patterns of IDC expression with spatial autocorrelations. To find the exact locations of IDC pressure within a field each year, IDC-susceptible varieties are planted in plots throughout the field, augmenting the varieties planted in incomplete blocks. Seed companies usually evalaute thousands of lines each year at each location, with at most two replicates; thus some lines may be planted in high and some in low IDC-pressure areas. The ephemeral nature of IDC spatial and temporal variation for high and low IDC pressure cannot be planned for, given the small plot evaluations of early-stage field trials, and thus require spatial models to correct for variation in IDC pressure within fields.

IDC phenotype scores typically range from 1 (the most resistant) to 5 (the most susceptible) in reports by academic field breeders [8], or from 1 to 9 by some commercial plant breeding organizations. In both cases, the scores represent ordinal data [9]. While plant breeders attempt to create ordinal data that forces IDC values to behave as continuous variables, the actual IDC scores often change in adjacent plots sharply from 1 to 9. This is consistent with what is observed in farmers fields. IDC symptoms in Iowa and southern Minnesota often appear to consist of oval shaped patches (Fig 1), due to location of soil moisture. In summary, ordinal data such as IDC creates phenotyping and selection challenges for plant breeders due to the genetic complexity of the trait, the scoring of IDC as ordinal data, and the use of small plots with 3 to 8 plants in early-stage evaluations in fields with ephemeral sharply delineated patches of IDC.

**Fig 1.**
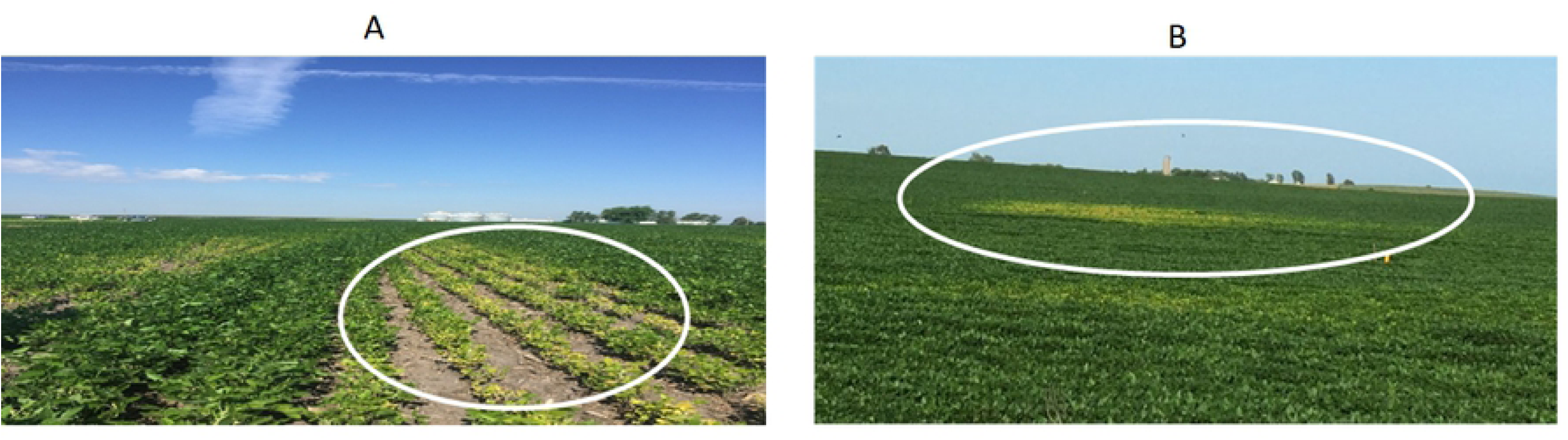
Spatial variation patterns in soybean IDC in commercial production fields. (A) IDC oval pattern in the lower/ditch area at Madrid, Iowa in 2016. (B) IDC circle pattern in a farmer’s field at Ames, Iowa 2016.

Accurate phenotypic data is the most important factor for both visual-based phenotype selection and marker-assisted selection. High-quality phenotypic data relies on both experimental design and accurate assessment of the phenotypes. Since presence of IDC is to some degree opportunistic, standard IDC resistant and susceptible checks or controls are included in field trials to estimate both the overall IDC pressure across the testing site and used as a reference to measure and adjust IDC scores of previously untested genotypes. To balance the dilemma of arrangement of check plots and maximize the number of test-line entries with field spatial variation, many experimental designs, such as augmented design [10], modified augmented design [11, 12], partial replicated [13], and augmented partially replicated (P-rep) designs [14-16] such as incomplete block alpha lattice [17, 18], have been developed to account for field plot variation. Also, different statistical models have been developed to account for field variation. Phenotype data quality control is the process of removing non-genetic variation caused by environmental noise from the estimated genotype values. Phenotypic data variation and subsequent patterns, or spatial variation, have been studied for many decades, especially in the geostatistics and econometrics discisplines [19-23]. Spatial models are used to account for autocorrelation among neighbors, which violates the identical independent distribution (iid) assumption for ordinary least square (OLS) analyses of variance [24]. Various spatial adjustment techniques have been developed to account for the spatial autocorrelation and significantly improve precision and repeatability of quantitative phenotypes. Collectively, these models can be clustered into three groups of spatial models based on the time of model development and the optimization mechanism used to adjust the spatial variation.

The first group includes the moving grid mean adjustment models [25-29]. The moving mean spatial analysis recently has become popular because the R package “mvngGrAd” implements the analysis. The package provides flexibility to pre-define any grid or pattern consisting of neighboring plots. The mean of the plots included in the grid is calculated and used as a covariate to account for the spatial variation [30]. In contrast with the spatial autoregressive (SAR) model, which treats spatial variation anisotropic along different directions [31], the moving mean average models treat spatial variation as isotropic, in which the covariates among neighbor plots are simple means of the neighbor plots within a user-defined grid. This approach has been reported to adjust the spatial variation successfully and thus has increased genomic selection accuracy from 0.231 to 0.37 for grain yield and from 0.436 to 0.614 for days to heading for wheat breeding projects [32].

The second and most extensively studied classification consists of the spatial autoregressive regression (SAR) models. These include parameters for autocorrelations among neighboring plots as covariates to model the correlated variation among neighbors. These models assme that closer the neighbors will be more highly correlated [33]. Versions of these models include spatial lag models, spatial error models, spatial lag plus error mixed models [34, 35], first-order autoregressive regression AR(1) or one-dimensional spatial analysis with row, column, or row + column [36], and the extended two-dimensional spatial analysis with interactions of rows and columns [37].

The SAR models focus on optimizing the variance-covariance structure of the residuals among neighbor plots. Evidence from a systematic comparison of covariance structures among experimental, spherical, Gaussian, linear, linear log, anisotropic power, and anisotropic exponential, show that AR(1) was generally not an optimal option for spatial analysis, and different covariance structures are needed to account for spatial variation at different trial sites [38], because each trial site has different variation patterns. Within this group of models, field variations are also divided into local trend or small-scale variation within a block or experiment, and global trend or large scale variation across the entire trial sites [39]. The nearest neighbor analysis model was developed to correct both local and global neighbors for the field variation [40, 41]. More complex models with polynomials were also developed to account for additive effects for either row or column, and non-additive epistasis interaction between the rows and columns [42]. To remove the local and global field variation effectively, Gilmour et al. [43, 44]proposed a sequential spatial model schema in which the first step removes local trend variation by fitting a two-dimensional range by row AR(1) and the second step is to remove the global trend variation by fitting one-dimensional polynomials or splines in the direction of rows or columns. To select the best model for each trial site, the sequential spatial model will run both model selection and model variable selection manually by applying graphical diagnostic tools to the spatial models. This manual model selection process was further extended with more optional models for the comparison and enhanced for model selection efficiency [45]. Over time the effectiveness of the spatial variation correction of SAR models have been improved. SAR analyses are routinely used for data analyses in geostatistics and econometrics [46].

The third group of models to account for spatial variation are known as “Tensor product penalized spline models,” or P-splines. These have been used to account for both local and global spatial variation in tree genetics using mixed models in Bayesian methods. Application of these methods to tree genetics increased the accuracy of estimated breeding values by 66% [47]. Bayesian mixed model methods were extended to 2-dimensional smoothed surface in an individual-tree based model using tensor product of linear, quadratic, and cubic splines for rows and columns, and the accuracy of breeding values for the offspring increased by 46.03% [48]. Most recently, an advanced P-spline model was proposed and developed as the R package Spatial Analysis of field Trials with Splines,”SpATS” [49]. This model includes both bilinear polynomial and smooth splines components: 1) The bilinear polynomial component consists of three sub-variables: row spatial trend, spatial column trend, and interaction between row and columns; and 2) The smooth spline component contains five smooth additive spatial components. This approach uses two-dimensional P-spline ANOVA representation of the anisotropic smooth surface formulated in a mixed model via SpATS. SpATS is advocated [49] because SpATS provides a one-step modeling approach by fitting a general SpATS model to analyze all field trials. The SpATS approach overcomes the sequential spatial model selection for different testing sites and can be used with large-scale, high-throughput and routine analyses of multiple environmental trials (METs). SpATS can fit both local and global variation, isotropic and anisotropic variation, one-dimensional and two-dimensional variation with one model, and optimize the best estimates of parameters to remove the noise from the true genotypic values without overfitting. With single step for model fitting, it minimizes the chance of using different models selected for different trials – which might lead to biases against different genotypes from different locations due to different selected model variables [50]. SpATS was tested by modeling spatial trends in sorghum breeding fields, and the results show that the improvement in precision and predicted genotypic values from SpATS analysis were equivalent to those obtained using the best SAR sequential model selection for each trial [50].

The three groups of spatial models for adjusting spatial variation have been mainly applied in econometrics and geostatistics while a few have been applied by plant breederes for continuous data, such as crop grain yield and plant height [28, 50, 51]. In contrast to continuous yield data, soybean IDC scores are discrete ordinal variables, whereas moving grid, SAR and P-spline models were developed and applied to continuous variables. In the applications, adjusted continuous trait phenotypes have improved precision and the repeatability of the field trials. To our knowledge, the application of these methods to adjust for non-genetic spatial patterns exhibited with ordinal traits such as IDC is limited to a single publication in which the moving grid was applied to IDC scores in soybean [28]. Also, while most of the published reports about spatial analysis models have investigated a few models, none have systematically compared the effectiveness of models for data obtained from fields with different levels of severity and irregular discontinuous spatial patterns. We hypothesize that the effectiveness of adjustments made by spatial models depends not only on the model parameters but also the severity of the spatial variation and the irregularity of variation patterns. The objective of research reported herein is: 1) to apply six different geospatial as well as two OLS models to three datasets with different levels of severity of spatial variation, variation patterns, field plot designs; and 2) to systematically compare models for spatial adjustment using R^2^, AIC, residual standard error, Moran’s I, and prediction accuracy.

## Materials and methods

A total of five data sets, four from field plot experiments, and one simulated provided a total of 11,602 unique genotypic lines (Table 1). Dataset 1 was simulated to mimic a circular IDC spatial variation pattern that can be found in fields in which potholes were drained with tiles in the last 80 years. The parameters used for the simulation are summarized in Table 2. Dataset 1 contains 1,050 simulated genotypes with a total of 2,100 IDC scores. The field layout consists of 42 ranges by 50 rows and genotypes are assigned to plots using a randomized complete block design with two replicates.

**Table 1.** Summary of the five data sets used to investigate application of geospatial models to remove non-genetic spatial patterns of plots evaluated for iron deficiency chlorosis.

| **Dataset Name** | **Exp. design** | **No. rows** | **No. range** | **No. entries** | **Ave No replicates** | **No. plots** | **Data sources** |
| --- | --- | --- | --- | --- | --- | --- | --- |
| Data set 1 | RCBD | 50 | 42 | 1,050 | 2.00 | 2,100 | simulated |
| Data set 2 | α-lattice | 220 | 26 | 2,774 | 1.83 | 5,074 | Iowa |
| Data set 3 | α-lattice | 24 | 220 | 2,652 | 1.79 | 4,740 | Minnesota |
| Data set 4 | α-lattice | 110 | 56 | 2,719 | 1.88 | 5,124 | Red River Valley |
| Data set 5 | α-lattice | 100 | 60 | 2,407 | 1.81 | 4,363 | Nebraska |
| Total |  | 504 | 404 | 11,602 | **<u>1.84</u>** | 21,401 |  |

**Table 2.** Summary of parameters used for data simulation. The first column is the list of variance components, columns 2, 3, and 4 are the variance values, distribution used for the simulation, and percentage of the variance components, respectively.

| **Variance Component** | **Variance** | **Distribution** | **Percentage of SD (%)** |
| --- | --- | --- | --- |
| Location SD | 1.5 | normal | 14.42 |
| Experiment SD | 0.5 | Poisson | 4.81 |
| Line SD | 2.1 | Normal | 20.19 |
| Range SD | 0.2 | Normal | 1.92 |
| Row SD | 1.0 | Normal | 9.62 |
| Rep SD | 0.1 | Normal | 0.96 |
| Pattern_SD | **3.8** | Normal | 36.54 |
| Residual | 1.2 | Normal | 11.54 |

In contrast to simulated dataset 1, datasets 2 to 5 represesnted experimental field data from 2016. They were selected to represent IDC-prone regions. For each set, the field plot design was a six-by-six alpha-lattice consisting of 36 plots, that were assigned 32 testing lines and four checks. These lines were in early stages of variety development projects, planted with two replicates in one location. The average number of replicates was 1.84 replicates/line across all the experiments. The reason for less than two replicates per testing line is due to low emergence rates at some sites.

IDC scores in data sets 2 to 5 were collected on individual hill plots from locations that have historically provided expression of IDC in soybeans. These sites were selected to minimize the IDC spatial variation by past years’ IDC records and the current year’s IDC status. Soybean IDC pressures vary from year to year for the same site, and testing sites do not always provide conditions for IDC expression every year. In order to assess potential IDC pressure at a site, several varieties previously determined to be IDC susceptible were planted about 10 days earlier than the expected planting date for all other lines. If the susceptible varieties showed IDC symptoms these standard IDC controls were then rogued and new experimental lines were planted at testing sites. Otherwise, these sites were not used as IDC testing sites.

IDC score scales from 1 for most tolerant to 9 for the most susceptible were used in all data sets. The IDC score criteria are 1 = green leaves (no chlorosis) to 9 = dead plant, following the rating scale presented in Fig 2. The IDC scores are treated as ordinal data.

**Fig 2.**
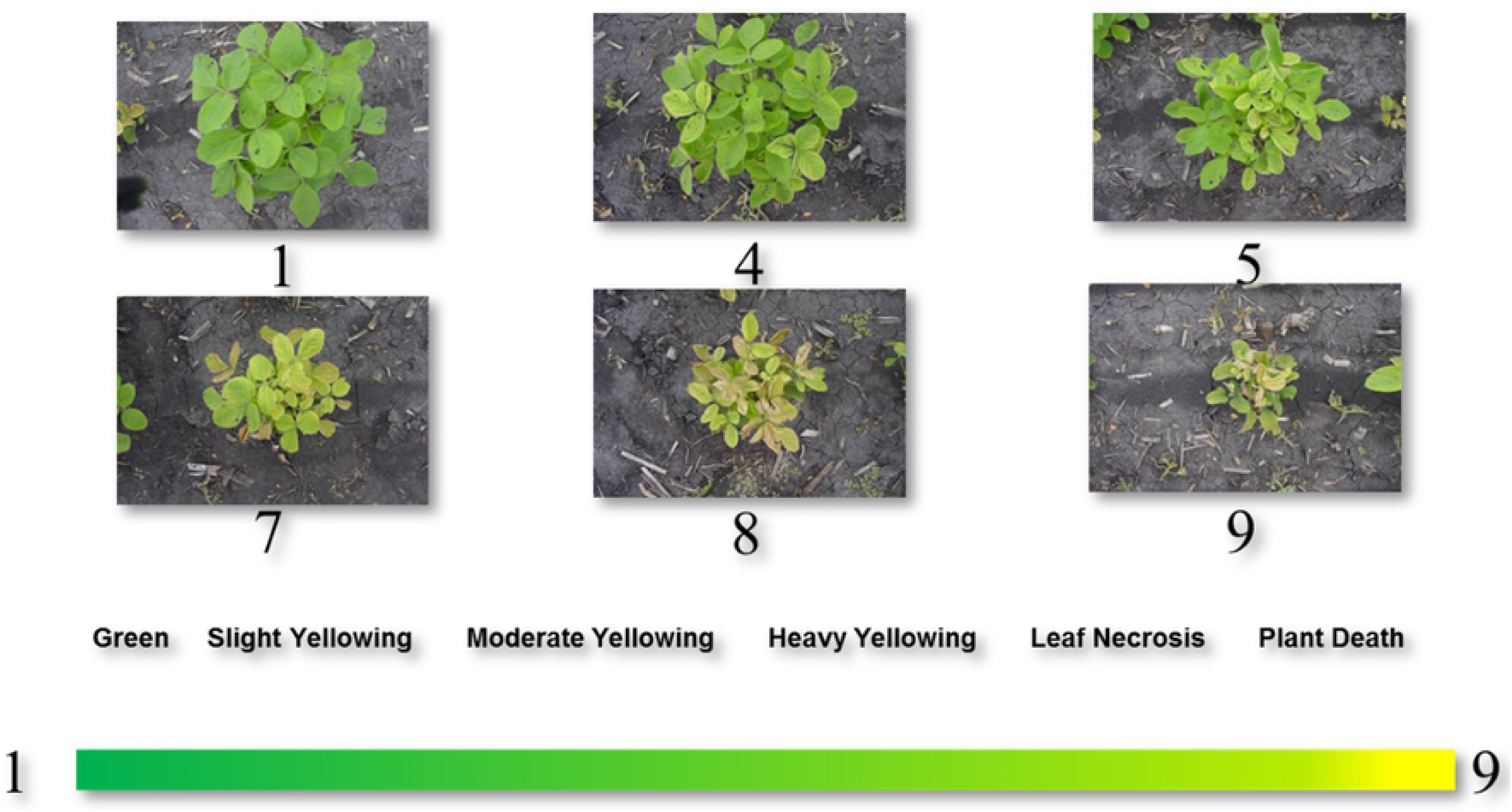
IDC score reference from 1 to 9 as ordinal data. 1 = green leaves (no chlorosis) to 9 = dead plant. Each image was taken from one hill plot with three to eight plants at growth stage V2 to V4.

Among the five data sets, we observed three classes of distinctive spatial variation patterns that we refer to as the Red River Valley (RRV) pattern, the Iowa and Minnesota (IA/MN) pattern, and the Kansas and Nebraska (KS/NE) pattern (Supplemental Fig 1). The three IDC field spatial patterns are consistent with the three soybean IDC-prone soil types, which were clustered by principal component analysis (PCA) based on 15 soil character measurements (Supplemental Fig 1). Five testing sites from North Dakota and Manitoba were clustered as the RRV group, four testing sites from Iowa and Minnesota were clustered together as the IA/MN group, and one testing site from Nebraska represented the KS/NE group (Supplemental Fig 1). KS/NE regions have relatively uniform IDC scores without noticeable spatial patterns, and no spatial model is needed to adjust plot IDC scores in this region.

The Red River Valey (RRV) IDC data has spatial variation among columns represented as block effects. In contrast, the IA/MN IDC testing sites show distinct spatial patterns relative to RRV and KS/NE regions. Thus, IDC data from the IA/MN region has spatial autocorrelation and needs spatial autoregressive analysis. Among the five testing sites from the IA/MN region, two show different spatial variation patterns (data sets 2 and 3) and are used to evaluate the geo spatial methods. All the results reported hereafter use these three data sets: two data sets from the IA/MN region, and one simulated data set (Table 1).

### Analytic methods

Including ordinary linear square (OLS), a total of eight models were compared (Table 3).

**Table 3.** Summary of the eight models compared in the research.

| Model Name | Model No | Spatial Term | R package | Reference |
| --- | --- | --- | --- | --- |
| OLS w/o RR | M1 | None | RMS | [52] |
| OLS w/ RR | M2 | None | RMS | [52] |
| MovingGrid | M3 | Mean of grid | mvngGrAd | [53] |
| SAR + lag | M4 | Lag | spdep | [54] |
| SAR + error | M5 | Error | spdep | [54] |
| SAR Durbin | M6 | Lag+ Error | spatialreg | [55] |
| ASReml AR1 | M7 | AR1(range): AR1(row) | ASReml-R | [56] |
| P-spline | M8 | psanova(range, row) | SpATS | [57] |

#### Model 1: Ordinary least square (OLS) without range and row covariates

OLS is a linear least-square method for estimating the unknown parameters in a linear regression model. OLS chooses the parameters of a linear function representing a set of explanatory variables by the principle of least squares. Traditionally the regression model does not include parameters for the spatial dependence of the experimental units. The general equation for the OLS with p variables can be written as:

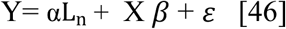

Where Y and *ε* are (n x 1) vector of the values of the response variable and the errors for the various observations, respectively. L_n_ is a (*n* x1) vector of ones associated with the constant term parameter α to be estimated. X is an (n×p) matrix of regressors or the design matrix. *β* is (1p x 1) vector of the parameters to be estimated.

OLS analysis for model 1 was conducted with the ols() function implemented the R package “regression modeling strategies (rms)” [52].

#### Model 2: Ordinary least square (OLS) with range and row

Model 2 analysis was similar to model 1 except two variables, range, and row were added in the model [45].

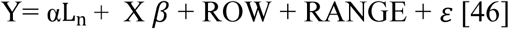

#### Model 3: Moving grid adjustment

The “moving grid average adjustment” is a spatial method to adjust for environmental variation in field trials. It is most common in field trials with few replicates, such as for early-stage breeding materials. All the raw data were aligned into a row by range rectangle layout. A grid is predefined based on the field variation pattern around each cell (= entry), and each observed value was adjusted by the values from the neighbor plots within the predefined grid. The mean of the cells included in the grid is calculated using the equation below [58]:

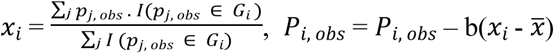

Where

*x*_*i*_ is the moving mean of the ith entry

*G*_*i*_ is the grid of entry i and I(·) is an indicator function that takes the value “1” if the condition is satisfied and “0” if not

*P*_*j, obs*_ are the observed phenotypic values of all entries which are included in *G*_*i*_

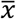 is the mean of all *x*_*i*_

b is the regression coefficient in the linear model.

*P*_*i, obs*_ are the adjusted phenotypic values of all entries.

The layout of the grid used to adjust the phenotype value is shown below in Fig 3:

**Fig 3.**
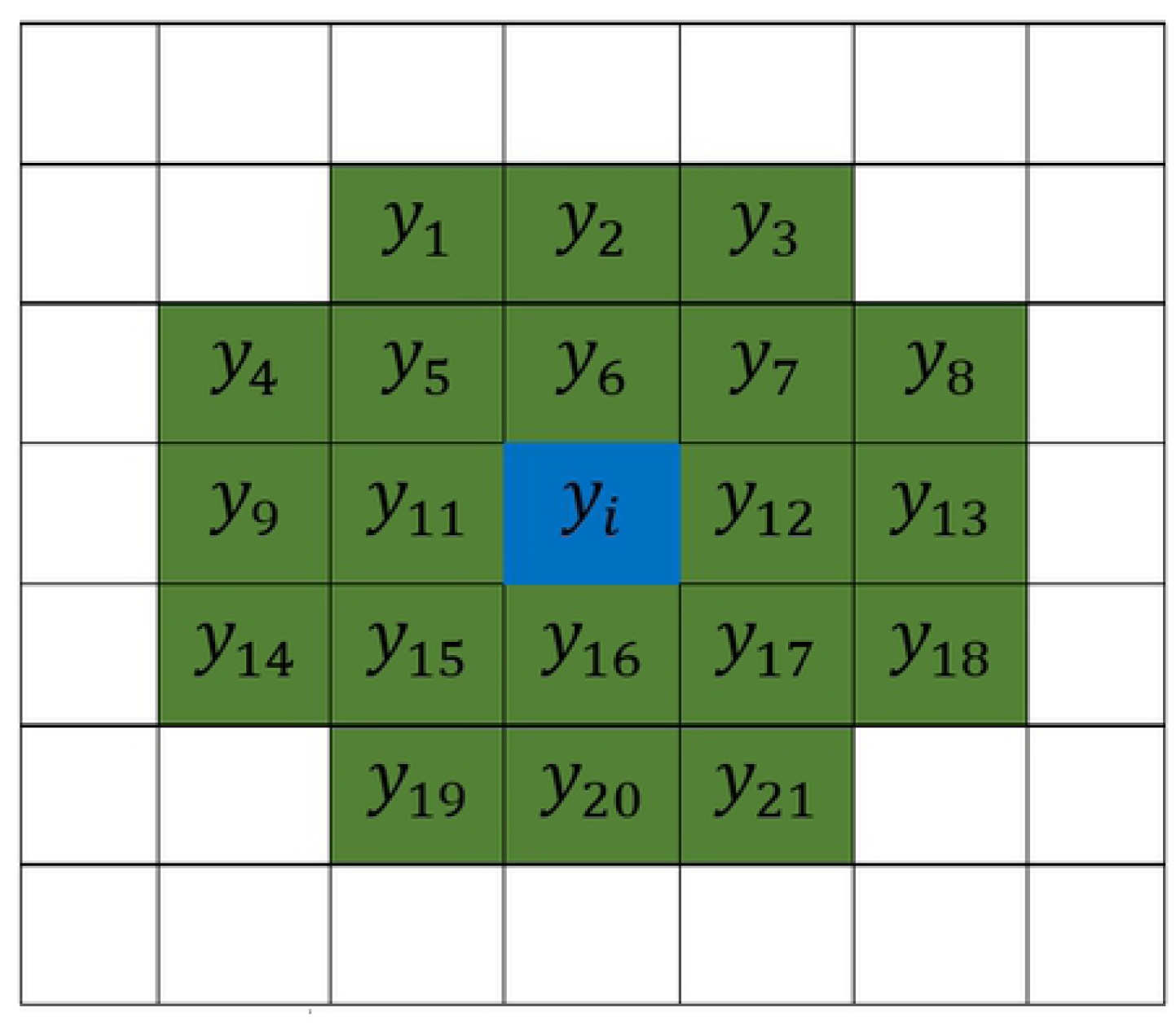
Layout of the grid used to calculate the mean.

The model executed for each data set was:

~~~
*movingGrid (rows = no.of.rows*,
      *columns = no.of.range*,
      *obsPhe=raw.observed.values*,
      *shapeCross=list(1:2,1:2,1:2,1:2)*,
      *layers=c(1:1)*,
      *excludeCenter = TRUE)*
~~~

where “shapeCross” is to set up the shape of the moving grid, “excludeCenter” is to define whether the center from each grid is included/excluded to calculate the mean.

#### Model 4: Spatial autoregressive lag model

When a value in one plot depends on the values of its neighbors, the errors are no longer uncorrelated and may not be independent and identically distributed (iid). Depending on the nature of the spatial dependence, OLS will be either inefficient, with incorrect standard errors, or will be biased and inconsistent [59]. When IDC scores depend not only on genetics, but also on scores from neighboring plants, the spatial lag model was introduced to correct for the autocorrelation [60]. In the spatial lag model, the spatial components were specified on the dependent variable, IDC scores. This setting leads to a spatial filtering of the variable, which are averaged over the surrounding neighborhood defined in W, called the spatially lagged variable. The spatial lag model can be specified as:

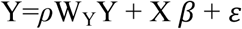

Where *ρ* is the autoregressive lag coefficient, which tells us how strong the resemblance is, on average, between Y_i_ and its neighbors; if *ρ* is not significantly different from 0, then spatial lag model becomes traditional OLS regression model.

*y*_*i*_ is the i^th^ IDC score, and *y*_*j*_ is all the neighbor’s IDC scores around i^th^ IDC score. *y*_*i*_ stands for one of n observed IDC scores, *y*_*j*_ stands for more than one IDC scores.

W_Y_ is a spatial weight matrix with n x p rows and ranges, describing the spatial correlation structure of the observations.

X is an n×p matrix of regressors or the design matrix; *β* is p x 1 vector of estimated coefficients.

Analysis of IDC scores with the spatial lag model was conducted via R package “spdep” [54] and “spatilreg” [55].

#### Model 5: Spatial autoregressive error model

In contrast to the spatial lag model treating autocorrelation as a lag component in the response variable, spatial error model regards the autocorrelation as part of the error term. The spatial error model incorporates a local and a spillover element in the variance-covariance matrix of the error term in a linear regression model [61]. Formally, the model can be written as:

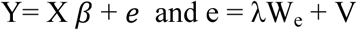

Where λ is the spatial error coefficient, if the absolute value of the λ is not significant bigger than 0, the spatial error model becomes the OLS regression model. W_e_ are the weight matrix to adjust the error correlation in the residuals.

Analysis of IDC scores with the spatial error model was conducted via R package “spdep”[54] and “spatilreg” [55].

#### Model 6: Spatial Durbin mixed model

A limitation of the spatial lag or spatial error models is that they can include either an autoregressive lag or a spatial error covariate in the model. In reality, some fields have complexed spatial variation with both autocorrelation lag and spatial error. Also, the dependencies in the spatial autoregressive relationships don’t only occur in the dependent variable but may also be present in the independent variables. The spatial Durbin mixed model was developed to account for dependent, the autocorrelation lag, spatial error, and independent variables [62, 63]. The model can be written as:

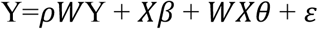

Where

Y, *X, ρ*, and *β* are defined as above

*W*Y: is the spatially lagged offering IDC scores accounting for various spatial dependencies with *W* defined as (n x n) spatial weight matrix

*ρW*Y: Endogenous interaction effect

*θ*: (k x 1) vector of unknown parameters

*θ WX*: Exogenous interaction effect

Implementation of the spatial Durbin model was carried out via the R package “spatilreg” [55]. *Model 7: AR1 by AR1 via ASReml-R*. The ASReml mixed model is widely used in plant and animal breeding and quantitative genetics. It also provides functions for spatial autoregressive analysis. ASReml-R is the only commercial R package in this study and used to compare whether it will outperform the free spatial analysis packages. One update in the new ASReml-R version 4 for spatial analysis, ASReml changed the random formula and error (rcov) component “rcov = ar1(range):ar1(row)” to residual = ar1(range):ar1(row) [56, 64].

The model used for the analysis is: asreml(fixed =IDC_scores ∼1, random = ∼ LINCD, residual = ∼ar1(range):ar1(row), data = IDC.data)

#### Model 8: P-spline mixed model implemented wth the R package SpATS

The P-spline approach models field trends using a smooth bivariate function of the range and row f(range, row), represented by a 2D P-splines [49, 65]. The P-spline method optimizes the fitted surface by penalizing the spatial effects. The degree of penalization over the fitted spatial variation trend is determined by smoothing parameters. The 2D range by row surface is decomposed into a sum of linear components and univariate and bivariate smooth function as:

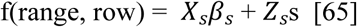

f(range, row) = *X*_*s*_*β*_*s*_ + *Z*_*s*_s [65] where:

*X*_*s*_*β*_*s*_ = *β*_*s*1_*row* + *β*_*s*2_*range* + *β*_*s*3_*row*. range

*Z*_*s*_s = *f*_1_(*row*) + *f*_2_(*range*) + *h*_3_(*row*).*range* + *h*_4_(*range*).row + *f*_5_(*row, range*)

*β*_*s*1_*row*: linear trend by row

*β*_*s*2_*range*: linear trend by range or column

*β*_*s*3_*row*. range: linear interaction trend by row x range

*f*_1_(*row*): main smooth trend across rows

*f*_2_(*range*): main smooth trends across ranges

*h*_3_(*row*).*range*: interaction trends between linear range by smooth surface row

*h*_4_(*range*).row: interaction trends between linear row by smooth surface range

*f*_5_(*row, range*): smooth-by smooth trends between ranges and rows

The P-splines mixed model can be written as:

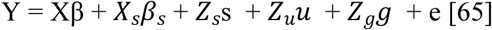

Where:

X, β, *X*_*s*_*β*_*s*_, *Z*_*s*_s are the same as above

*Z*_*u*_*u*: u is the sub-vector of random row and range effects accounting for discontinuous field variation; Zu = [*Z*_*row*_|*Z*_*range*_] is the design matrix

*Z*_*g*_*g*: g is the vector of random effects of genotypic effects of the testing lines or hybrids; *Z*_*g*_ are the design matrix for the genotype effects.

The final SpATS model R scripts used in this study for P-spline is as below:

~~~
P-Spline.model =
SpATS(response=“IDC_Scores”, genotype=“LINCD”, genotype.as.random = TRUE,
            spatial = ∼PSANOVA(row, range, nseg = c(10,10),
            degree = 3, nest.div = 2), fixed = NULL,
            control = list(tolerance = 1e-03, monitoring =1),
            data = IDC.data)
~~~

The term “spatial” is an auxiliary function used for modeling the spatial or environmental effect as a two-dimensional penalized tensor-product of marginal B-spline basis functions with anisotropic penalties based on the PSANOVA approach [66, 67]. Inside spatial, “nseg” stands for the number of segments in the P-splines, 10 and 10 segments were used for both range and row, respectively. Parameter “degree” stands for numerical order of the polynomial of the B-spline basis for each marginal. Degree of 3, cubic B-splines was used for the IDC analysis. Parameter “nest.div” is a divisor of the number of segments (nseg) to be used for the construction of the nested B-spline basis for the smooth-by-smooth interaction component. In this case, the nested B-spline basis will be constructed assuming a total of nseg/nest.div segments. The value was set to 2 for the IDC analysis.

### Performance metrics

The effectiveness of models were compared using: 1) coefficient of determination, R^2^; 2) Akaike’s information criterion (AIC); 3) residual standard error (RSE); 4) Moran’s I index; 5) p-value of Moran’s I; 6) prediction accuracy I (whole data set); 7) and prediction accuracy II (cross-prediction accuracy).

All except Moran’s Index are broadly used metrics for many statistical methods. Moran’s index also known as the spatial autocorrelation index simultaneously measures spatial autocorrelation based on both feature locations and feature values [69]. With a set of features and an associated attribute, Moran’s I index evaluates whether the pattern expressed is clustered, dispersed, or random. Moran’s I test provides a way to check whether there is spatial autocorrelation in the field data and whether residuals from a spatial model are not correlated or randomly distributed, with the iid property. Moran’s I test value is between −1 to +1. A value of “-1” indicates the large and small values intersperse across the field and the data are negatively auto-correlated, while “+1” indicates high IDC scores surrounded by high IDC scores or low IDC scores surrounded by low IDC score; these scores are positively auto-correlated. If all the residuals from a model are iid, and there is no autocorrelation, then Moran’s I should be close to zero. If I is close to 0, then the spatial adjustment is successful. If I is close to either −1 or +1, with p-values <0.05, then interpretation is that the adjustment by the model does not remove a significant amount of the spatial autocorrelation

Software that implements these models do not generate estimates of all of these metrics, thus the metrics were calculated manually:

1. *R*^*2*^: *R*^*2*^*= 1-*SSresidual SStotal [68] where *SS*_*residual*_ is the sum of squares of the residual from the model, and *SS*_*total*_ is the total sum of squares from the data.
2. *AIC: AIC = 2k-2ln(L) = −2(log-likelihood) + 2K* [68] Where K is the number of model parameters (the number of variables in the model plus the intercept). Log-likelihood is a measure of model fit. The higher the number, the better the fit. For AIC, the smallerthe value, the better fit of the model.
3. Residual standard error (RSE) = square.root (MSE) where MSE is mean square error.
4. Moran’s I index: Moran’s I index was calculated as:

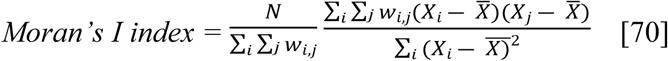

Moran’s I index was calculated via *moran.test* from the R package “spdep,” as below (order of the data is important using function “moran.test.” The data need to be the same order as the one in the weight matrix of the rectangle):

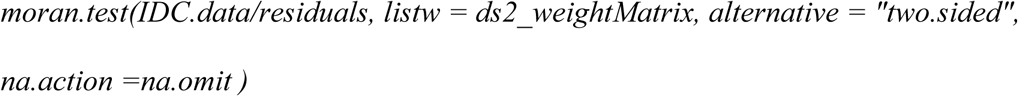

Where N is the number of spatial units indexed by i and j; X is IDC score; 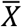 is the average IDC scroe; *w*_*i,j*_ is the spatial weight matrix..
5. P-value of Moran’s I: The null hypothesis for the Moran’s I test in that there is no autocorrelation among the data in the area, and the data collected are randomly distributed. If the p-value from Moran’s I test is not significant, or the p-value > 0.05, the spatial distribution of feature values may be the result of random spatial processes.
6. Prediction accuracy I (whole data set): is calculated as

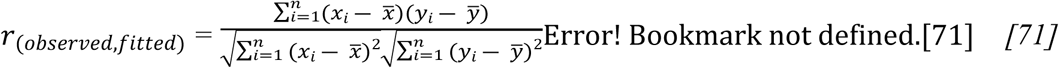

Where n is the number data points or sample size *x*_*i*_ from 1 to n is the observed values *y*_*i*_ from 1 to n is the fitted values from model
7. Prediction accuracy II (cross prediction): is the correlation coefficient between the overserved and predicted for the testing lines planted in low IDC pressure area. Formula for prediction accuracy II is the same as prediction accuracy I except using the testing lines planted in the low IDC pressure regions.

### Comparison methods

#### Heatmap and Lagrange Multiplier Test

Heatmaps of IDC field layout were made using the python package “seaborn” and the R package “fields” [72]. The Lagrange Multiplier Test was conducted to select the best model among all the spatial autoregressive (SAR) models via function “lm.LMtests “from the R “spdep” package.

#### Relative efficiency (RE)

The mean squares error (MSE) from each analysis were used to estimate the relative efficiency (RE) of each model relative to the OLS models:

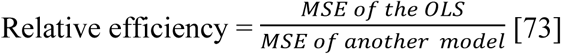

## Results and discussion

### Assessments of data set complexity

Spatial effective dimension is a measure of the complexity, in which larger EDs are indicative of more complex patterns [49]. The shapes of evident patchy spatial patterns of the IDC scores were best explained by the integration of one-dimensional range or row trends (functional trend row or range and surface range or row) and two-dimensional trends (interaction function trend row by range “Row: Range,” surface trend range by linear function trend row “F(Range): Row,” linear functional trend range by surface trend row “Range: F(Row),” and surface trend interaction range by row “F(Range): F(Row)” (Table 4). Overall, one-dimension surface range trends were more complicated than that of row, in which all the f(range) > f(row). For data set 1 f(range) 3.0 > f(row)=2.3, for data set 2 the values are 7.0 > 6.6, and for data set 3 5.8 > 3.3 (Table 4). From the two-dimension level, 2D surface trend rows by surface trend ranges “f(row):f(range)” are the most significant spatial terms. The percentage of the effective spatial dimensions are 58.42, 44.12, and 53.92% for data sets 1, 2, and 3, respectively. Another observation is that linear trend for row plays a large role (13.9 in data set 2) and linear trend range did not contribute at all (0 in data set 2). Opposite results were obtained from data set 3: 24.6 for the linear trend of range and 17.6 for the linear trend for row. The differences of linear trend ranges and rows between data sets 2 and 3 indicate the spatial structures of data set 2 and 3 are different.

**Table 4.** Summary of effective spatial dimensions (EDs) of the smooth surface components for the three data sets.

| Variables Name | Data Set 1 |  | Data Set 2 |  | Data Set 3 |  |
| --- | --- | --- | --- | --- | --- | --- |
|  | ED<sub>s</sub> | ED<sub>m</sub> | ED<sub>s</sub> | ED<sub>m</sub> | ED<sub>s</sub> | ED<sub>m</sub> |
| Range | 10.1 | 42 | 0.0 | 220 | 24.6 | 221 |
| Row | 41.1 | 50 | 13.9 | 28 | 17.6 | 24 |
| Row:Range | 1.0 | 1 | 1.0 | 1 | 1.0 | 1 |
| F(Range) | 3.0 | 11 | 7.0 | 11 | 5.8 | 11 |
| F(Row) | 2.3 | 11 | 6.6 | 11 | 3.3 | 11 |
| F(Range):Row | 0.0 | 11 | 6.5 | 11 | 5.8 | 11 |
| Range:F(Row) | 1.8 | 11 | 1.6 | 11 | 10.0 | 11 |
| F(Range):F(Row) | 83.3 | 121 | 28.9 | 36 | 79.7 | 121 |
| % F(Range):F(Row) | 58.42 | 46.90 | 44.12 | 10.94 | 53.92 | 29.44 |
| Total | 142.6 | 258 | 65.5 | 329 | 147.8 | 411 |

Results of variance components analysis of the three random (LINCD, Range, and Row), five surface variables (f(RANGE), f(ROW), f(RANGE): ROW, RANGE:f(ROW)), and one 2D surface range trend by surface row trend, f(RANGE):f(ROW)) are summarized in Table 5. These show that: 1) three datasets have large differences in level of spatial variances, with dataset 1 having the smallest total variance of 431.5 (Table 5) and dataset 2 having the largest variance of 6597.2 (15-fold larger than that of dataset 1); 2) surface range trend by surface row trend f(RANGE):f(ROW), accounts for the majority of the variance in dataset 1 and 2 – up to 85.6% and 89.6%, respectively; and 3) dataset 3 has different spatial variation patterns from datasets 1 and 2; a smaller variance component from surface range trend by surface row trend of 15.14%. In contrast, linear range trend by surface row trend, range:f(row), takes up to 78.1% in the dataset 3. One observation is that variance components for LINCDs are very small: 0.1%, 0.01%, and 0.05% for datasets 1, 2, and 3, respectively. The overall mean-variance components for LINCDs is only 0.05% across the three data sets. Similar results were observed for the residual variance, in which the mean percentage of variance component is only 0.1% across three datasets (last column in Table 5). The overall results from the variance components analysis show that spatial variation along the rows are much larger than that from ranges. The mean percentages related to surface row term “f(ROW)” are 5.95% for “f(ROW),” 26.42% for “RANGE:f(ROW),” and 63.44% for “f(RANGE):f(ROW)” with a total of 95.81% (last column in Table 5).

**Table 5.** Summary of variance component analysis from the tensor product panelized P-splines. Var stands for variance, type for data type: “R” for random and “S” for surface variable.

| Variables | Type | Data set 1 |  | Data set 2 |  | Data set 3 |  | Mean |
| --- | --- | --- | --- | --- | --- | --- | --- | --- |
|  |  | Var | % var | Var | % var | Var | % var | % var |
| LINCD | R | 0.43 | 0.10 | 0.71 | 0.01 | 0.46 | 0.05 | 0.05 |
| Range | R | 0.01 | 0.00 | 0.00 | 0.00 | 0.02 | 0.00 | 0.00 |
| Row | R | 0.07 | 0.02 | 0.20 | 0.00 | 0.78 | 0.08 | 0.03 |
| f(RANGE) | S | 4.97 | 1.15 | 39.05 | 0.59 | 13.09 | 1.39 | 1.04 |
| f(ROW) | S | 55.63 | 12.89 | 223.32 | 3.39 | 14.92 | 1.58 | 5.95 |
| f(RANGE):ROW | S | 0.31 | 0.07 | 352.16 | 5.34 | 32.58 | 3.45 | 2.95 |
| RANGE:f(ROW) | S | 0.55 | 0.13 | 66.26 | 1.00 | 737.68 | 78.13 | 26.42 |
| f(RANGE):f(ROW) | S | 369.15 | 85.56 | 5912.83 | 89.63 | 142.97 | 15.14 | 63.44 |
| Residual | R | 0.34 | 0.08 | 2.61 | 0.04 | 1.67 | 0.18 | 0.10 |
| total |  | 431.47 | 100.00 | 6597.15 | 100.00 | 944.18 | 100.00 | 100.00 |

### Comparisons of models

The relative rankings for the models based on different evaluation metrics indicates each model has advantages and disadvantages for correcting spatial patterns in the IDC data. A summary of performance metrics for the eight models applied to data set 1 in Table 6 suggests that the P-spline model implemented with SpATS provides the largest R^2^ of 0.8931, prediction accuracy of 0.9473, and residual standard error of 0.5582. In contrast, the moving grid adjustment is the least desired model because it has the smallest R^2^ (0.3409), lowest prediction accuracy (0.6246), and the largest Moran’s I index (0.9492). For data set 1, model OLS with range and row is the second-best based on prediction accuracy (0.9200), which is much better than OLS without range and row in the model indicating that row and range terms are important for correcting the spatial variation associated with rows and ranges of the grid. Another observation is that ASReml has the second largest R^2^ (0.7289) and second smallest residual standard error (0.6364), but prediction accuracy (0.8538) is low and ranked the sixth among the eight models. If Moran’s I index is used to rank the models, SAR + mixed model is the best since it has the smallest Moran’s I index (0.0349) and the largest p-value of the Moran’s I index (0.02179). For all eight models, estimates of prediction accuracy II is lower than that of prediction accuracy I indicating that the testing materials planted in high IDC pressure areas have higher prediction accuracy than those planted in low or no IDC pressure areas. AIC values are widely disparate among the models. For example, only two parameters are used for the moving grid average, while the P-spline, requires many more parameters:142.6 for data set 1 and 147.8 for data set 3 (Table 4) to fine-tune the boundary of the spatial trends and thus has a much larger AIC than the other models.

**Table 6.**
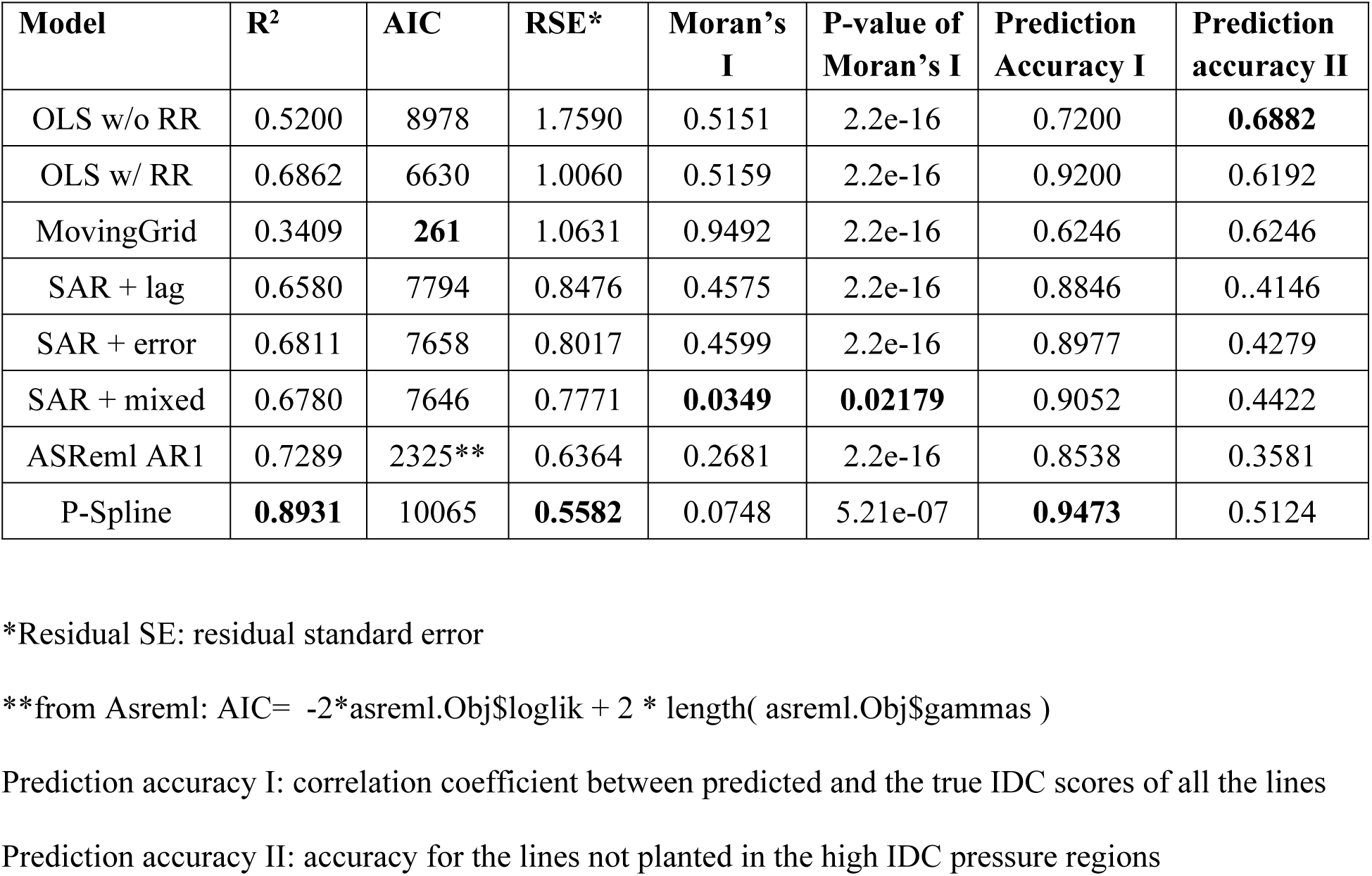
Summary of the comparison among different models via R^2^, residual variance, prediction accuracy, Moran’s I index from data set 1. Highlighted are the most desirable values in each column.

| Model | $R^2$ | AIC | RSE* | Moran's I | P-value of Moran's I | Prediction Accuracy I | Prediction accuracy II |
| --- | --- | --- | --- | --- | --- | --- | --- |
| OLS w/o RR | 0.5200 | 8978 | 1.7590 | 0.5151 | 2.2e-16 | 0.7200 | **0.6882** |
| OLS w/ RR | 0.6862 | 6630 | 1.0060 | 0.5159 | 2.2e-16 | 0.9200 | 0.6192 |
| MovingGrid | 0.3409 | **261** | 1.0631 | 0.9492 | 2.2e-16 | 0.6246 | 0.6246 |
| SAR + lag | 0.6580 | 7794 | 0.8476 | 0.4575 | 2.2e-16 | 0.8846 | 0.4146 |
| SAR + error | 0.6811 | 7658 | 0.8017 | 0.4599 | 2.2e-16 | 0.8977 | 0.4279 |
| SAR + mixed | 0.6780 | 7646 | 0.7771 | **0.0349** | **0.02179** | 0.9052 | 0.4422 |
| ASReml AR1 | 0.7289 | 2325** | 0.6364 | 0.2681 | 2.2e-16 | 0.8538 | 0.3581 |
| P-Spline | **0.8931** | 10065 | **0.5582** | 0.0748 | 5.21e-07 | **0.9473** | 0.5124 |

\*Residual SE: residual standard error
\*\*from Asreml: $AIC = -2 * asreml.Obj\$loglik + 2 * length( asreml.Obj\$gammas )$
Prediction accuracy I: correlation coefficient between predicted and the true IDC scores of all the lines
Prediction accuracy II: accuracy for the lines not planted in the high IDC pressure regions

The results from Table 6 are generally consistent with the eight heatmaps of the residual plots from data set 1 (Fig 4). The residual heatmap from model 1, 2, 4, 5, and 6 show a “baseball” pattern, indicating little improvement in autocorrelations among the plots, whereas models 3, 7, and 8 show a circle pattern, indicating better adjustments to remove autocorrelations among the residuals. The heatmap from using the P-Spline model provides the greatest number of plots with random residuals, and its legend bar has the smallest scales (from −2 to 1), whereas the rest of the models have scales ranging from from −2 to 2 or 3. The residual heatmap from “MovingGrid” is the typical case for positive autocorrelation close to 1, where all the residuals inside the circle are −1 values of +3 outside the circle. Moran’s I index of the residual heatmap from “MovingGrid” model also is the largest and is close to 1. Based on estimated performance metrics in Table 6 and Fig. 4, it appears that the best models for adjusting plot IDC scores in data set 1 are either the “SAR + mixed” and “P-Spline via SpATS,” and the least desirable model is the “MovingGrid”.

**Fig 4.**
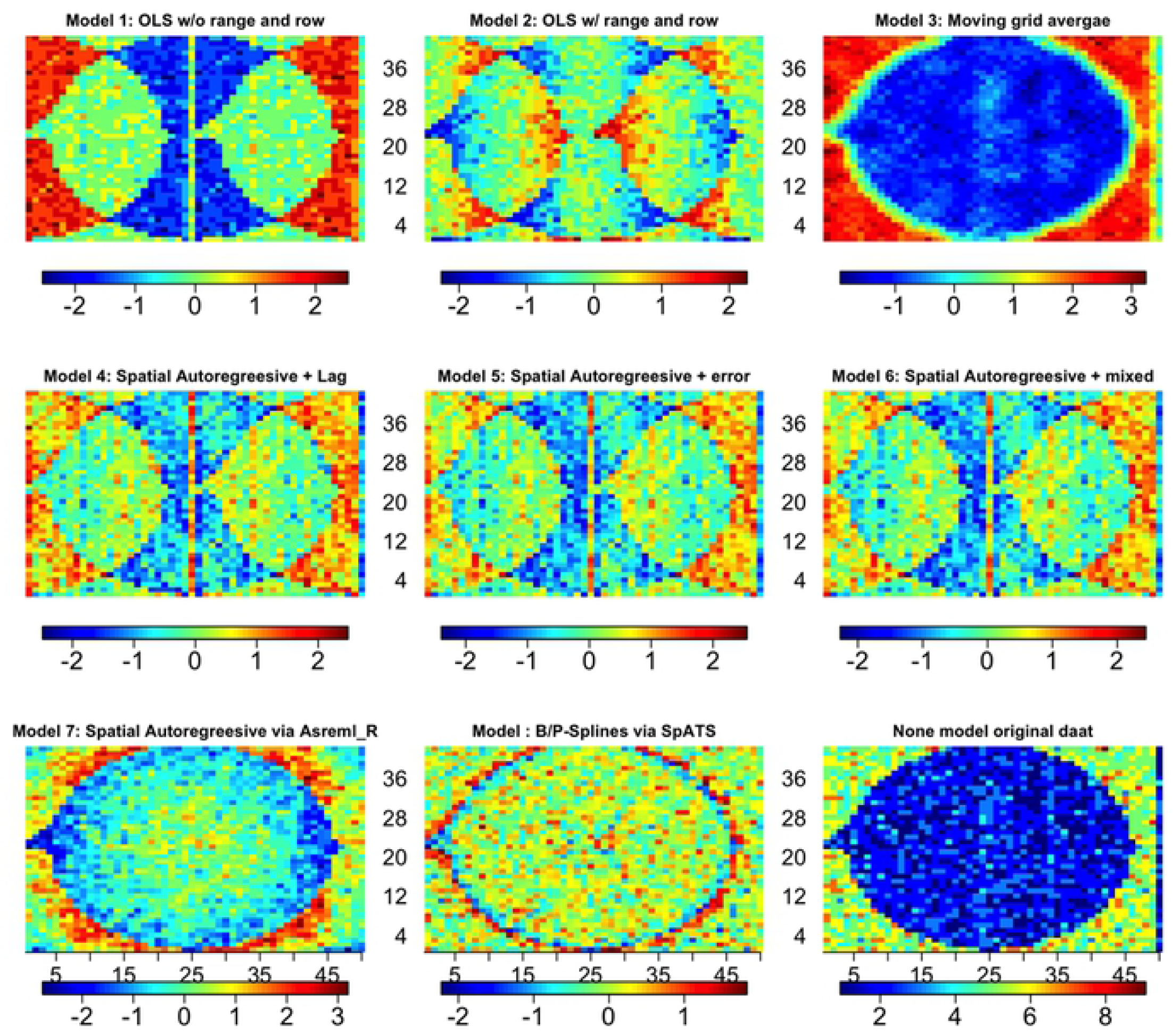
Heatmaps of residual values from eight models applied to data set 1. In each of the figures, x-axis is the rows from 1 to 50; y-axis is the ranges 1 to 42. The numbers under legend bars are the range of residual values, and the numbers above the legend bar in the bottom half of the figure are the row numbers.

Performance metrics and heatmaps of residuals for the eight models applied to data set 2 are summarized in Table 7 and Fig. 5. From the highlighted values in Table 7, the group of SAR models with either lag, error, or mixed have the best performance metrics among the eight models. Based on R^2^, residual standard error (RSE), and Moran’s I index, SAR + error term as a covariate in the model ranks the best. If based on prediction accuracy, Model SAR + mixed is the best since it has the highest prediction values: 0.9486 and 0.8435 for prediction accuracy I and II, respectively. In contrast to the results from data set 1, the model AR1⊗ AR1 via ASReml has the second worst R^2^and it is only slightly better than that of the “MovingGrid” model.

**Table 7.** Summary of the comparison among different models via R^2^, residual variance, prediction accuracy, Moran’s I index from data set 2. OLS w/o RR stands for ordinary least square without range and row; OLS w/ RR for ordinary least square with range and row. Highlighted are the most desirable values in each column.

| Model | $R^2$ | AIC | RSE | Moran's I | P-value Moran's I | Prediction Accuracy 1 | Prediction Accuracy 2 |
| --- | --- | --- | --- | --- | --- | --- | --- |
| OLS w/o RR | 0.666 | 22657 | 1.937 | 0.2148 | 2.2e-16 | 0.9428 | 0.8295 |
| OLS w/ RR | 0.648 | 22793 | 1.923 | 0.2131 | 2.2e-16 | 0.9429 | 0.8283 |
| MovingGrid | 0.385 | **6788** | 1.8094 | 0.8664 | 2.2e-16 | 0.5839 | 0.7426 |
| SAR + lag | 0.685 | 22521 | 1.3458 | 0.1655 | 2.2e-16 | 0.8356 | 0.5540 |
| SAR + error | **0.715** | 22479 | **1.3080** | **0.1518** | 2.2e-16 | 0.8457 | 0.5752 |
| SAR + mixed | 0.701 | 22581 | 1.3408 | 0.1778 | 2.2e-16 | **0.9486** | **0.8435** |
| ASReml AR1 | 0.555 | 14689 | 1.6765 | 0.3962 | 2.2e-16 | 0.8088 | 0.5033 |
| P-Spline | 0.565 | 50678 | 1.6156 | 0.2639 | 2.2e-16 | 0.7636 | 0.4636 |

**Fig 5.**
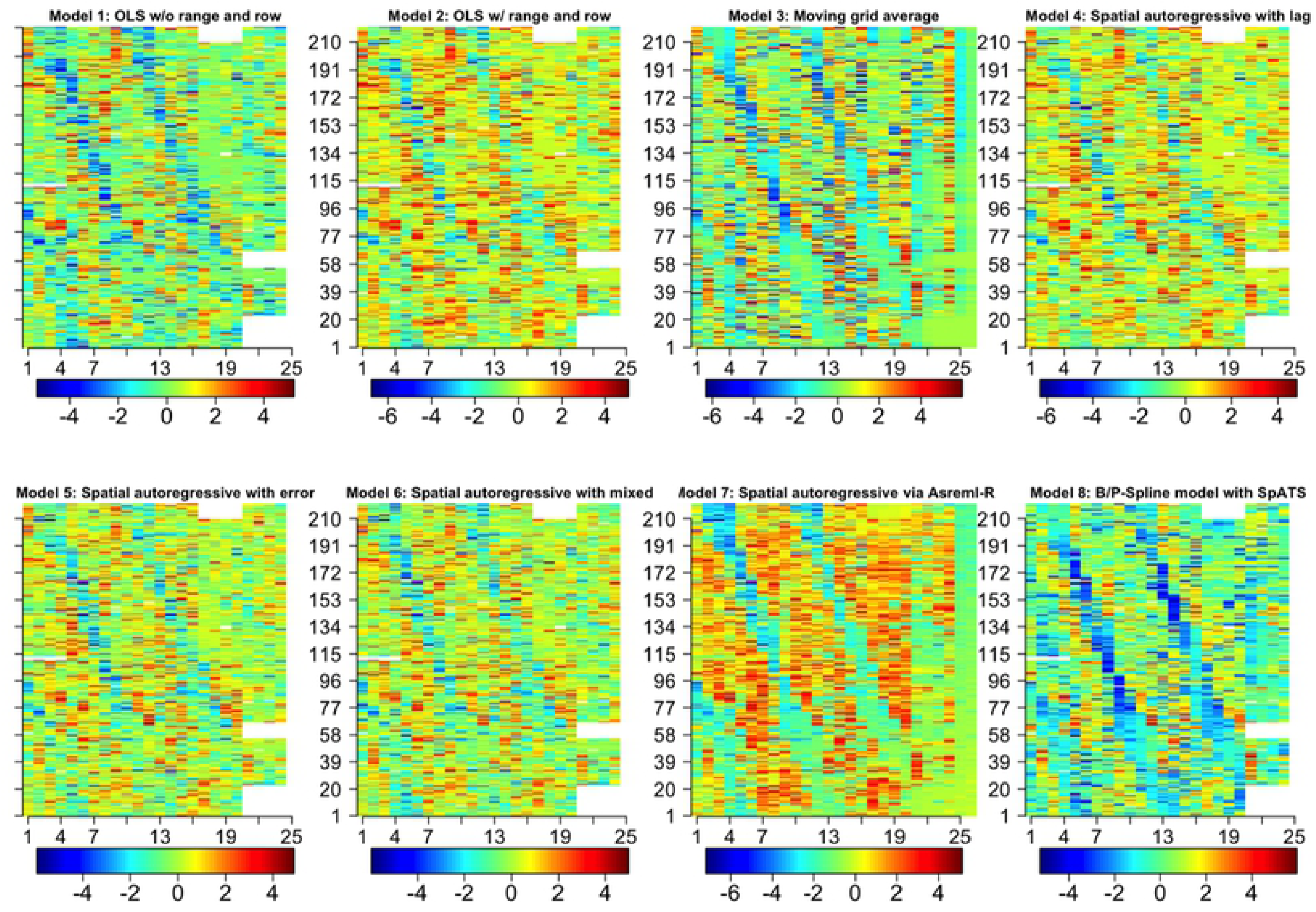
Heatmaps of residuals derived for each of eight models applied to data set 2. Each of the images has 220 ranges by 26 rows. X-axis is row and Y-axis is the range. The numbers under legend bars are the range of residual values, and the numbers above the legend bar in the bottom half of the figure are the row numbers.

In contrast to application of models to data set 1, the performance metrics from the P-spline model applied to data set 2 were not the best, most of the best performance metrics were from the SAR models. Based on AIC values the P-spline model has the largest AIC value 50,678, suggesting that the P-spline model is overfitting the data. Three models, “OLS w/o RR,” “OLS w/ RR,” and “SAR + mixed”, have very similar prediction accuracies, 0.9428, 0.9429, and 0.9486, respectively, indicating that range and row effects are not as important for making adjustements in data set 2 as for data set 1. This result is consistent with the total spatial effective dimensions (ED_s_) in Table 4, in which data set 2 has a total of only 65.5 ED_s_ with a large field of 220 range by 26 rows, whereas data set 1 has an ED_s_ of 142.6 with a small field of 50 ranges by 42 rows. Heatmaps from the eight models applied to data set 2 show clear spatial variation patterns (Fig 5), indicating that none of the methods fully remove the spatial patterns.

The performance metrics from application of the eight models to data set 3 are very similar (Table 8) to results for data set 2. The results of the three SAR models, “SAR + lag,” “SAR + error,” and “SAR + mixed,” are similar. The model “SAR + mixed” has the highest R^2^ (0.9491), prediction accuracy (0.9746), and smallest RSE (0.5827) (Table 8). However, in contrast to the results from data set 2 “SAR + lag” instead of model “SAR + error” has the smallest Moran’s index (0.022).

**Table 8.** Summary of the comparison results among the eight models via R^2^, AIC, residual standard errors, prediction accuracy, Moran’s I index from data set 3. Highlighted are the most desirable values in each column.

| Model | $R^2$ | AIC | RSE | Moran’s I | P-value of Moran’s I | Prediction Accuracy 1 | Prediction Accuracy 2 |
| --- | --- | --- | --- | --- | --- | --- | --- |
| OLS w/o RR | 0.8390 | 20298 | 1.8440 | 0.1456 | 2.2e-16 | 0.9534 | 0.8020 |
| OLS w/ RR | 0.8442 | 20145 | 1.8150 | 0.1358 | 2.2e-16 | 0.9541 | 0.8033 |
| MovingGrid | 0.5685 | **5204** | 1.6790 | 0.9306 | 2.2e-16 | 0.6569 | 0.4049 |
| SAR + lag | 0.9076 | 18251 | 0.7852 | **0.0220** | **0.2330** | 0.9672 | 0.8471 |
| SAR + error | 0.9450 | 17283 | 0.6059 | 0.0568 | 4.5e-08 | 0.9723 | 0.8654 |
| SAR + mixed | **0.9491** | 17180 | **0.5827** | 0.0826 | 1.8e-15 | **0.9746** | **0.8753** |
| ASReml AR1 | 0.6399 | 12603 | 1.5503 | 0.4233 | 2.2e-16 | 0.8448 | 0.5004 |
| B/P-Spline | 0.7494 | 41241 | 1.2935 | 0.1169 | 2.2e-16 | 0.8684 | 0.5732 |

Heatmaps of the residuals from application of the eight models to data set 3 (Fig 6) indicate that models 3 and 7 produce patterns that are very similar to the heatmap of the raw IDC scores, while models 4, 5, 6, and 8 retain, albeit limited, spatial patterns. Comparing the legend scales, the models “SAR + error” and “SAR + mixed,” have the smallest residual ranges from −3 to 5, and all the others have the residual ranges from −4 to +4 except that adjustments from the “MovingGrid” with residual ranges from −2 to +4. The heatmap of residuals from application of model “SAR + lag,” appears to have the least evidence of a spatial pattern, and Moran’s I is close to 0.

**Fig 6.**
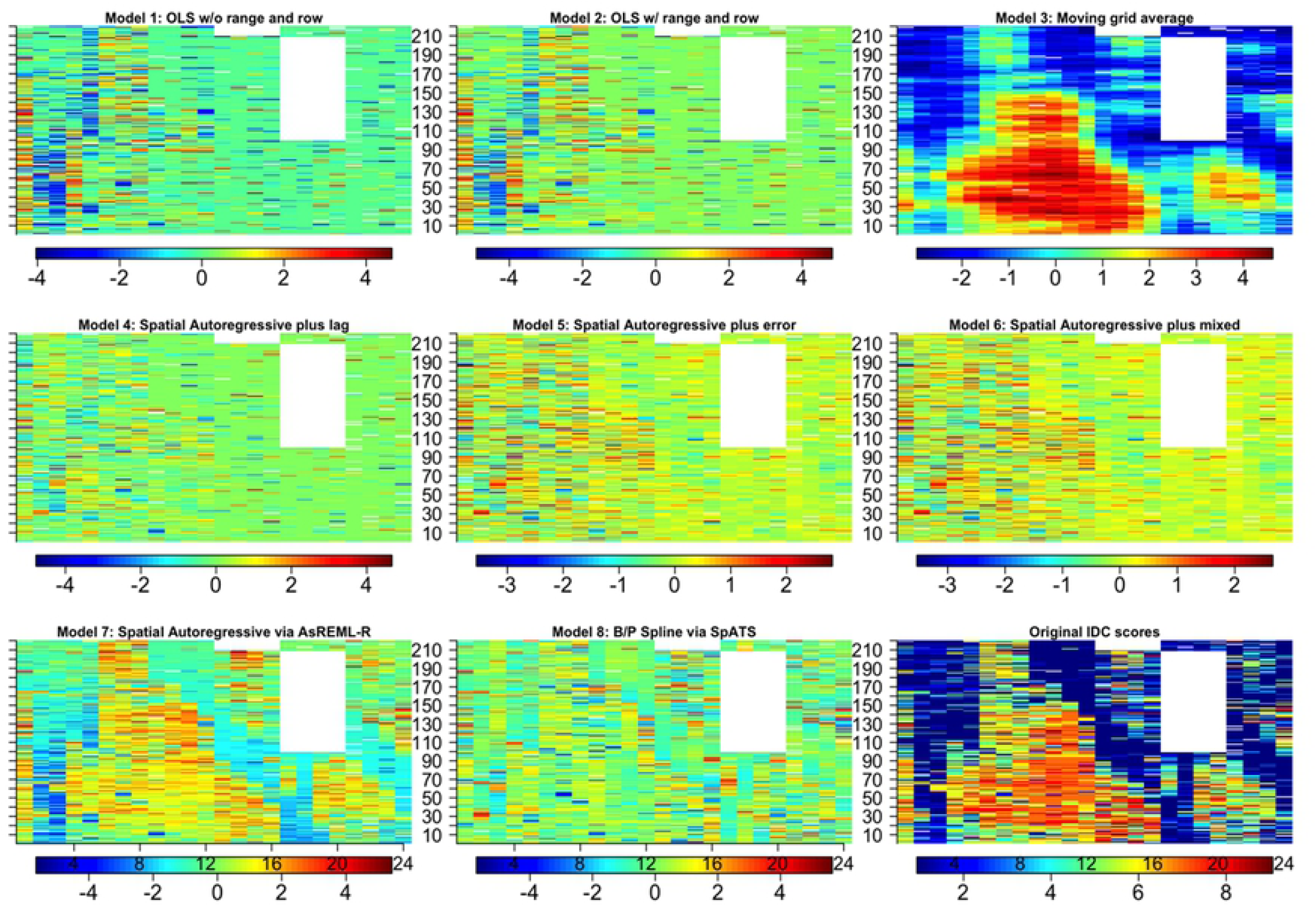
Heatmaps of residuals from data set 3 derived from eight models applied to IDC scores. The numbers under legend bars are the range of residual values, and the numbers above the legend bar in the bottom half of the figure are the row numbers.

Based on the results in Tables 6, 7, and 8 and Figures 4, 5, and 6, application of “SAR + mixed,” seems to provide the best outcomes for data sets 2 and 3, and model 8, “P-Spline via SpATS,” provided the best adjustements to IDC scores for data set 1.

With limited knowledge of the distributions of RSE associated with each model from the three data sets, we chose to conduct a distribution-free nonparametric Kruskal-Wallis test to assess whether there is a significant difference among RSE values generated by the eight models (Table 9). Model 1, “OLS w/o RR “was the worst, and Model 1, 2, and 3 are not significantly different. Similarly, model 6, “SAR + mixed” is the best, but models 4, 5, 6, 7, and 8 are not significantly different. As shown in Fig 7, three groups were observed with models 1, 2, and 3 in group I, models 4, 5, and 6 in group II, and Models 7 and 8 in group III. These three groups are consistent with the model’s mathematical and spatial covariate structures. Pairwise t-tests among the three groups show that group I and II are significantly different, with a p-value < 0.001, Groups II and III show significant differences with p-value < 0.01. Group I and III are statistically different at p-value of 0.1, but not 0.05 (Table 10).

**Table 9.**
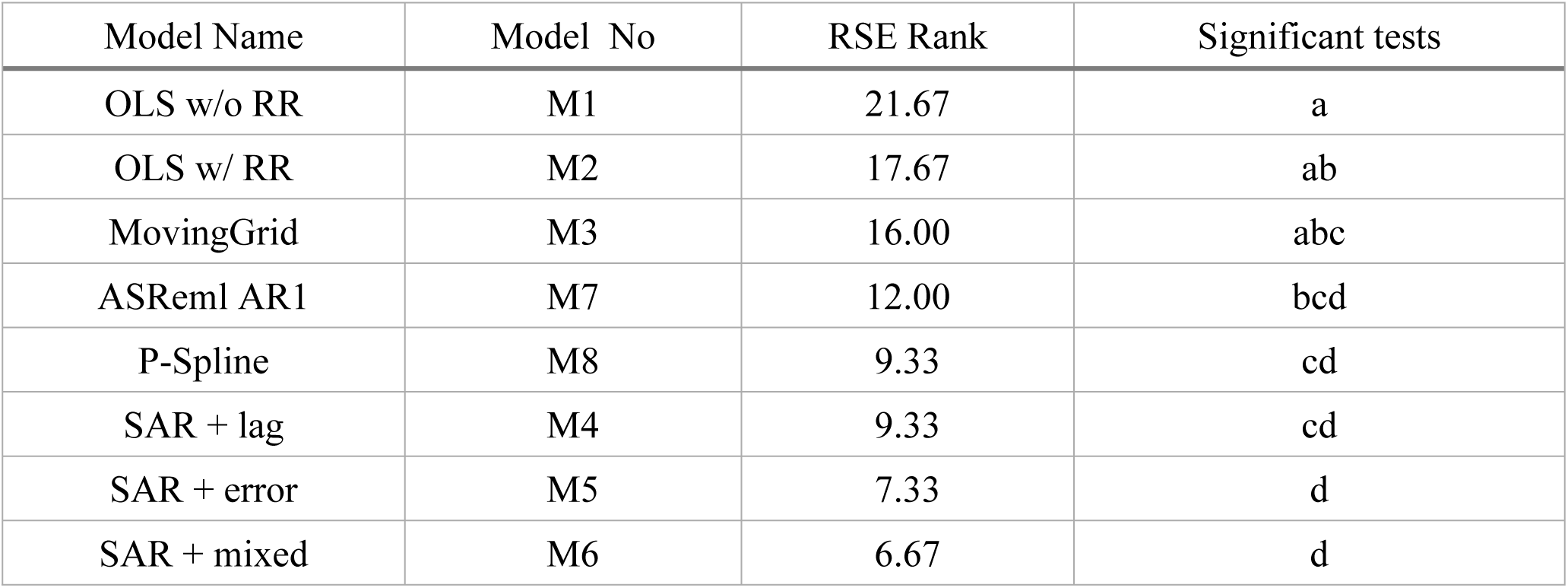
Results of the Kruskal-Wallis test.

**Table 10.** Pairwise t-test of three groups of spatial models.

| Model group | Mean of 1<sup>st</sup> group | Mean of 2<sup>nd</sup> group | Mean difference | t-test | p-values |
| --- | --- | --- | --- | --- | --- |
| Groups I vs II | 1.78 | 0.66 | 1.12 | -6.3367 | 0.00039 |
| Groups I vs III | 1.78 | 1.42 | 0.36 | -2.0133 | 0.08396 |
| Groups II vs III | 0.66 | 1.42 | -0.76 | 3.5051 | 0.00993 |

**Fig 7.**
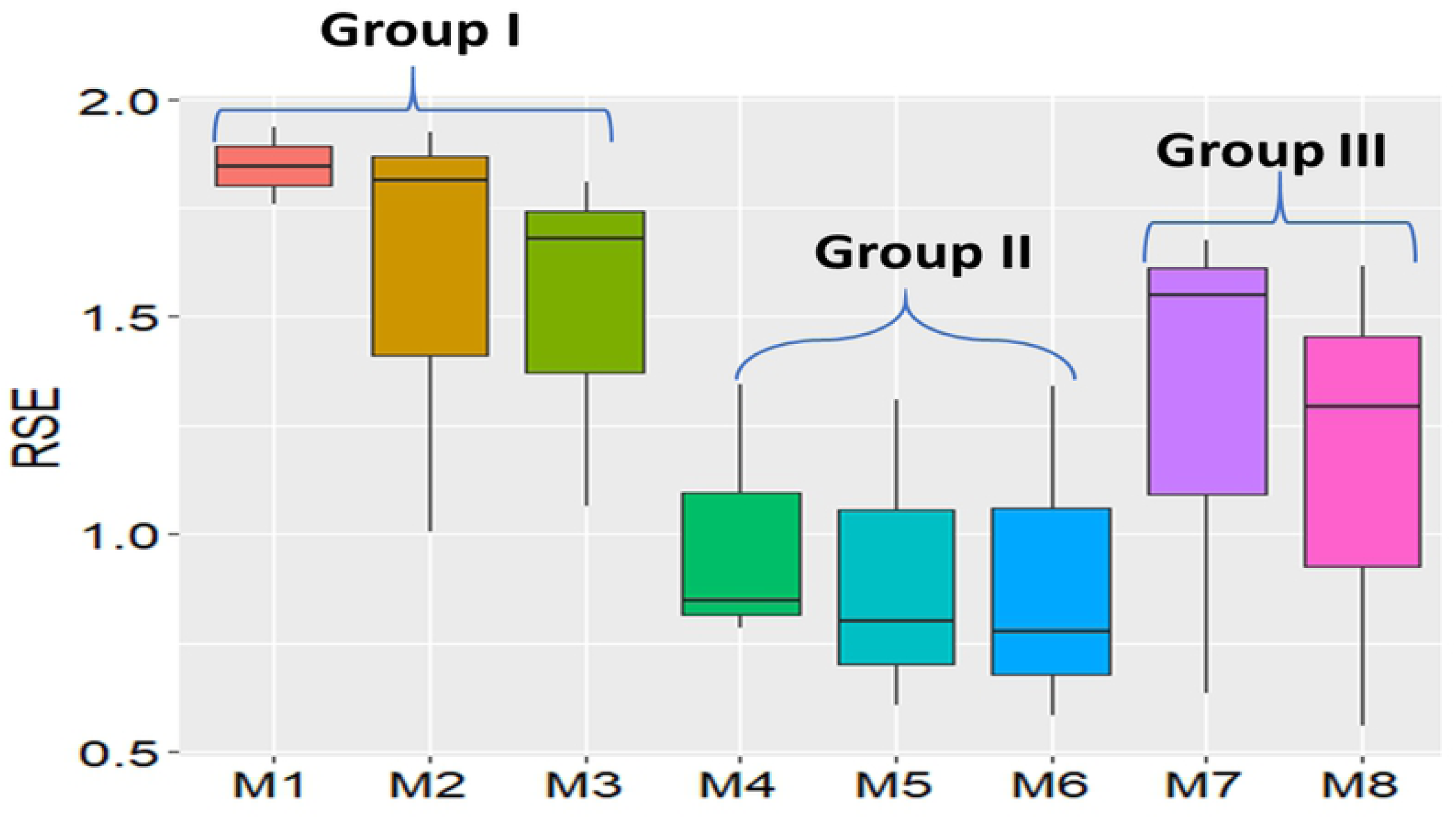
Boxplot of the residual standard error (RSE) generated by application of eight spatial models to three data sets consisting of range x row organization of plots scored for IDC. Group I consists of OLS and moving grid models, group II consists of SAR models and Group III consists of ASReml AR1 and P-spline models.

### Relative Efficiency (RE) of the spatial autoregressive (SAR) analyses

Relative to the ordinary least square models without (OLS w/o RR) and with range and row information (OLS w RR) efficiencies of models are presented in Table 11. The model “SAR + mixed” has the highest relative efficiency of 4.21indicating that at least four additional replicates of the OLS w/o range and row information would be needed in order to attain the same residual error. All relative efficiencies from the models are larger than 1.0, demonstrating that range and row information will explain significantly greater amounts of variance among IDC scores from all three data sets. The last column of Table 11 shows the relative efficiency of the seven models compared with Model 2, ordinary least square (OLS) with range and row in the model (OLS w/ RR). All the six models with range and row, have relative efficiencies larger than 1.0. The relative efficiencies of these six models indicates that there are significant improvements to be realized using more than simple “range” and “row” information to correct for spatial autocorrelation for spatial variation of IDC scores.

**Table 11.** Summary of relative efficiency (RE) of spatial analysis of all models compared to. ordinary least square (OLS) without range and rowand ordinary least square (OLS) with range and row.

| **Model Name** | **RE to OLS**<br>**w/o RR** | **RE to**<br>**OLS w/ RR** |
| --- | --- | --- |
| OLS w/o RR | 1.00 | 0.86 |
| OLS w/ RR | 1.36 | 1.00 |
| MovingGrid | 1.48 | 1.04 |
| ASReml AR1 | 2.06 | 1.23 |
| SAR + lag | 3.46 | 1.59 |
| B/P-Spline | 2.55 | 1.37 |
| SAR + error | 4.16 | 1.75 |
| SAR + mixed | 4.21 | 1.76 |

### Lagrange Multiplier Test (LMT)

Based on performance metrics and relative efficiencies the Group II models provided the best outcomes from adjusting IDC scores collected from real field trials (data sets 2 and 3). The Lagrange Multiplier Test (LMT) provides an overview of spatial dependence by these types of models (Table 12). In order to choose within this group of models larger dependence estimates are preferred. From the dependence estimates, spatial lag and spatial error models have very similar dependence estimates of 2728.1 and 2730.3, respectively. Tests for possible presence of lagged variance except Lmerr, RLMerr, is 108.98. Similarly, tests for possible presence of error variance (except LMlag, RLMlag), is 106.75. The last test is for both error and lag model, “SAR+mixed”, its dependence estimate is 2837.1 which is larger. Based on LMT test results, “SAR+mixed” should be used to correct for autocorrelations. However, much greater computation time is needed for SAR models. For data set 2, it took >10 hours to run the “SAR + mixed” model with 5,720 data points and a total of ∼25 hours to run the three data sets via a desktop Windows 10 with 16 Gb RAM and a core i7 CPU. The P-splines model implemented in the SpATS R package took less than 15 minutes for all three datasets.

### Effects of field plot operations on spatial analyses

The basic principles of experimental design are randomization, replication, and local control [74]. For IDC evaluations, replication and local control with an incomplete alpha-lattice design were used in data sets 2 and 3. But, randomization of previously untested lines is seldom practiced. Breeders usually group lines by families to ease visual evaluations. For example, a typical soybean breeding program might evaluate 32 lines per family from 400 families for a total of 12,800 lines. The IDC trials will be grouped into 400 trials associated with each of the families. The lines within a family will be randomly assigned to plots within each trial. But the 400 trials representing variability among families are usually not randomized across the field site. Rather, the families are arranged by pedigree and relative maturity for purposes of visual comparisons and to accommodate operational activities such as avoiding inter-plot competition between early and late maturing lines and avoiding damage to plots from mechanical harvest. A harvest combine and other equipment will begin with plots that are planted with early maturing families and proceed through the field from the earliest to latest maturing families. If a family is created by crossing resistant by resistant lines or moderately resistant by moderately susceptible lines, there will be patches of resistant or susceptible lines associated with families and these will create patterns unrelated to IDC. The genotypic patterns may confound the non-genetic spatial variation pattern. Under this circumstance, the spatial analysis may lead to biased adjustments. There is an opportunity to develop methods that will include covariates for genotypic relationships among families and expected maturities of families in the spatial models.

Plot size may affect spatial patterns and subsequently the effectiveness of spatial autoregressive models. Soybean IDC field trials were planted in hill plots, not row-plots (Fig 8). Each hill plot contained eight seeds. Hill plots were spaced 15 inches from center to center between ranges and 10 inches from center to center between rows (left-hand image in Fig 8). In contrast plots used to evaluate continuous variables such as yield and plant height are usually much larger (right-hand image in Fig 8). Plot size has been shown to impact the effectiveness of spatial models [75]. Larger plot sizes were better for spatial adjustements of yield values grown in heterogeneous fields. However, contrary to results indicating large plot sizes are better, evidence from a 28-year case study for optimizing experimental designs show that the relative efficiency of the Randomized Complete Block Design increased by 240% as the length of two-row plots decreased from 5.6 m to 1.4 m [76]. Similar results from a comparison of different spatial models among correlated error, nearest neighbor analysis, and autoregressive regression AR(1) indicates that smaller plot size is more efficient to capture spatial variation and thus increase the relative efficiency of the experimental design [77]. Both small plot size and ordinal IDC scores may be related to the high residual standard error we found in application of the P-spline model to data sets 2 and 3.

**Fig 8.**
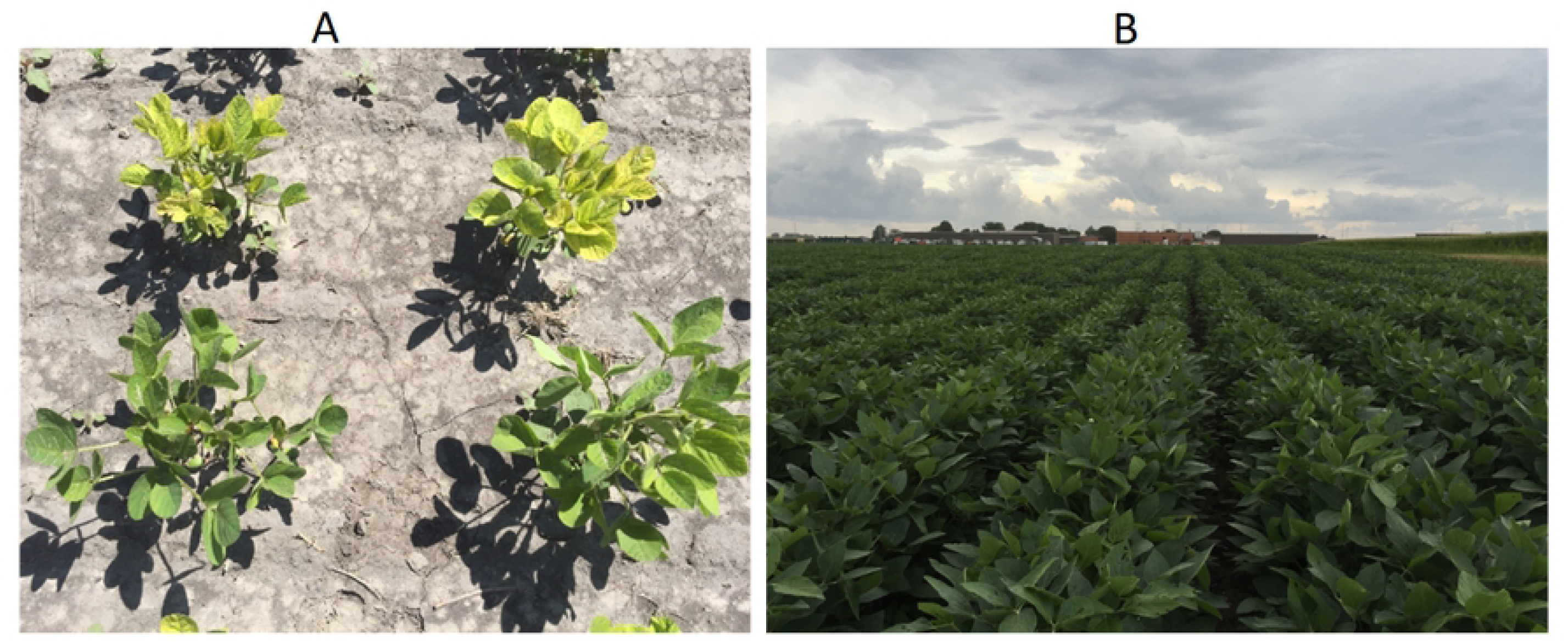
Images of IDC hill plots vs. yield test plots. Left image shows chlorosis phenotype of two susceptible testing lines (top two hill plots) and a two resistant testing lines (bottom two hill plots). Right image shows yield test plots with plants in 10m rows.

### Inconsistent results relative to previous field plot studies

Previously, when applied to continuous traits, P-spline models have been shown to be more effective and efficient than SAR models [49, 50]. For our three data sets application of the P-spline model was best only when applied to data set 1. For this simulated data set the P-splie model identified the discontinuity boundary between iron deficient and non-deficient conditions (Fig 9, bottom left-hand heatmap). Using a mixed model to obtain best linear unbiased predictions of the IDC values for each of the lines, we observed a ‘baseball’ pattern in the heatmap (Fig 9, bottom middle). Recall that the underlying genotypic values were randomly simulated based on a normal distribution. Moran’s index of the residual plots is 0.0748 with p-value 5.21e-07, which was much smaller than the threshold p-value 0.05. Comparing the Moran’s index, 0.5152 from the raw IDC data, p-value less than 2.2e-16, the spatial autocorrelation coefficient is dramatically reduced, from 0.5152 to 0.0748, for the P-spline model. But, in terms of goodness-of-fit, there is still autocorrelation left in the residual plots. There is a noticeable circle in the residual plot (Fig 9, top right-hand) demonstrating that even this best model did not remove the spatial autocorrelation pattern completely.

**Fig 9.**
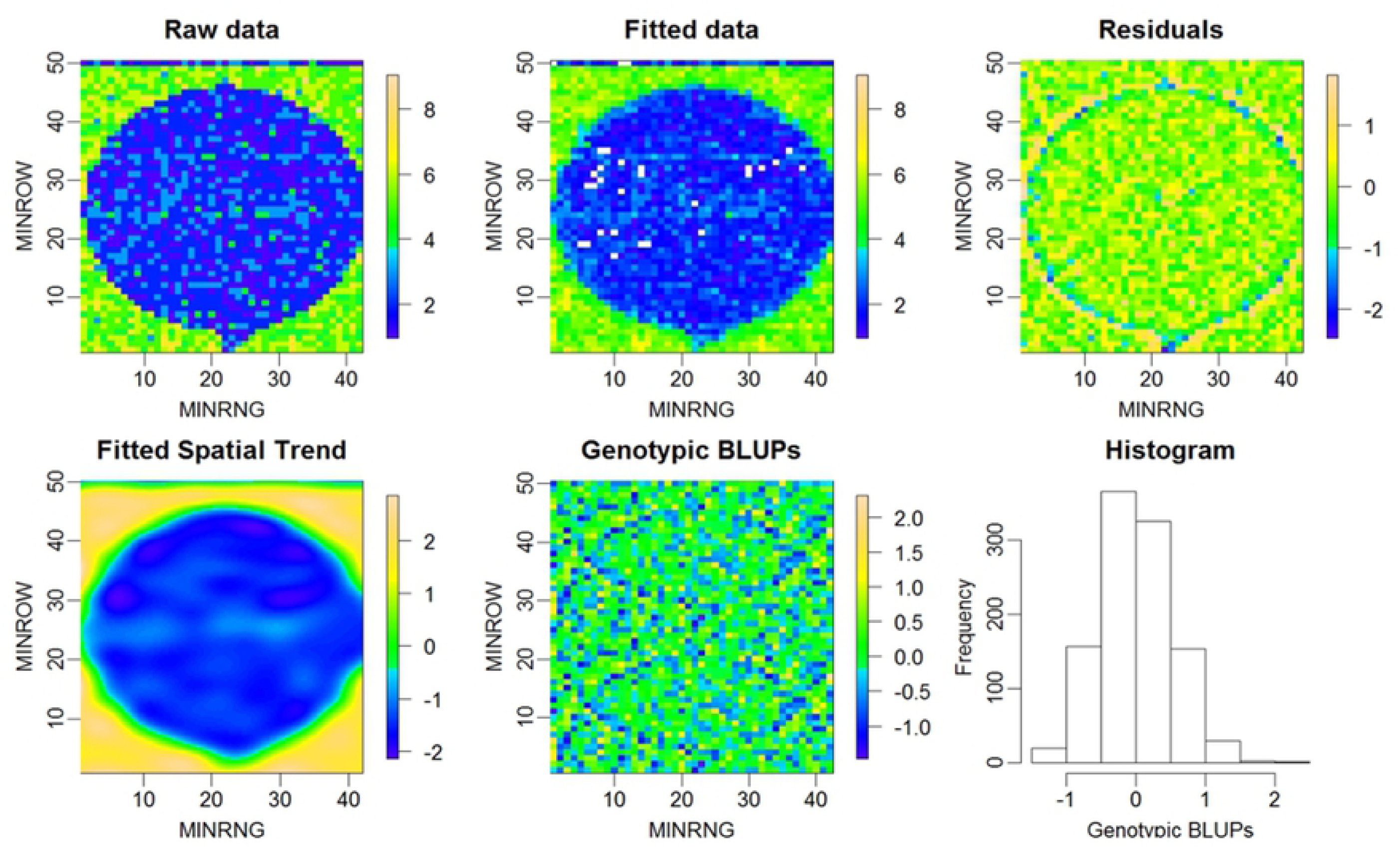
Heatmaps and histogram from application of the P-spline model to data set 1. The plots are organized in 42 rows (X-axis) by 50 ranges (Y-axis) with a total of 2,100 plots. Genotypic BLUPs are the best linear unbiased predictions of breeding values for the simulated genotypic IDC values and range from −2 to +2.

Application of the P-spline model to data set 2 indicate a different spatial pattern than that from the simulated data set 1 (Fig 10). Note that the heatmap of the fitted trend uses a continuous-like grid, which was smoothed by 2D P-splines. The spatial surfaces displayed irregular patchy patterns across field 2, and the discrepancy of the spatial trends between raw and fitted pattern remain after adjustments by all eight evaluated models (Fig 5). The heat map of the residual plots has a very similar spatial pattern as both fitted values and raw IDC score data, demonstrating that there is still significant spatial autocorrelation after application of the P-spline model (Fig 10).

**Fig 10.**
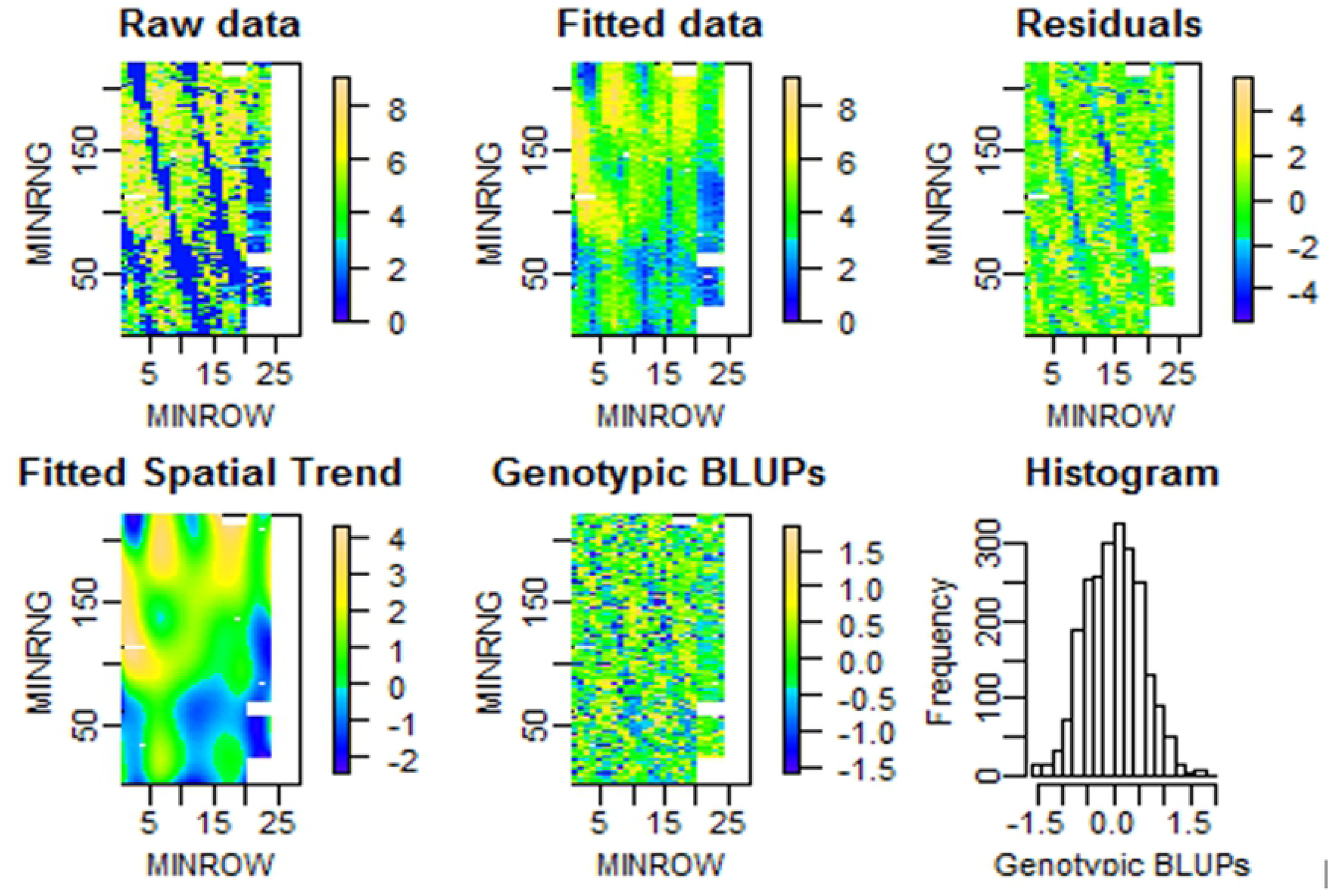
Heatmaps and histogram from data set 2. The rectangle layout consists of 26 rows (X-axis) by 220 ranges (Y-axis) and a total of 5,720 plots. The white area in the right side of each heatmap are plots without recorded data.

The P-spline model performed better for data set 3 (Fig 11). The fitted spatial trend matched the raw field pattern very well, and heatmaps of both genotypic BLUPs and residuals appear to represent randomly distributed variables. No clear noticeable spatial pattern exists like the results obtaind from application to data sets 1 and 2, even though the abolute value of the Moran’s I index is still greater than 0 (p-value < 0.05).

**Fig 11.**
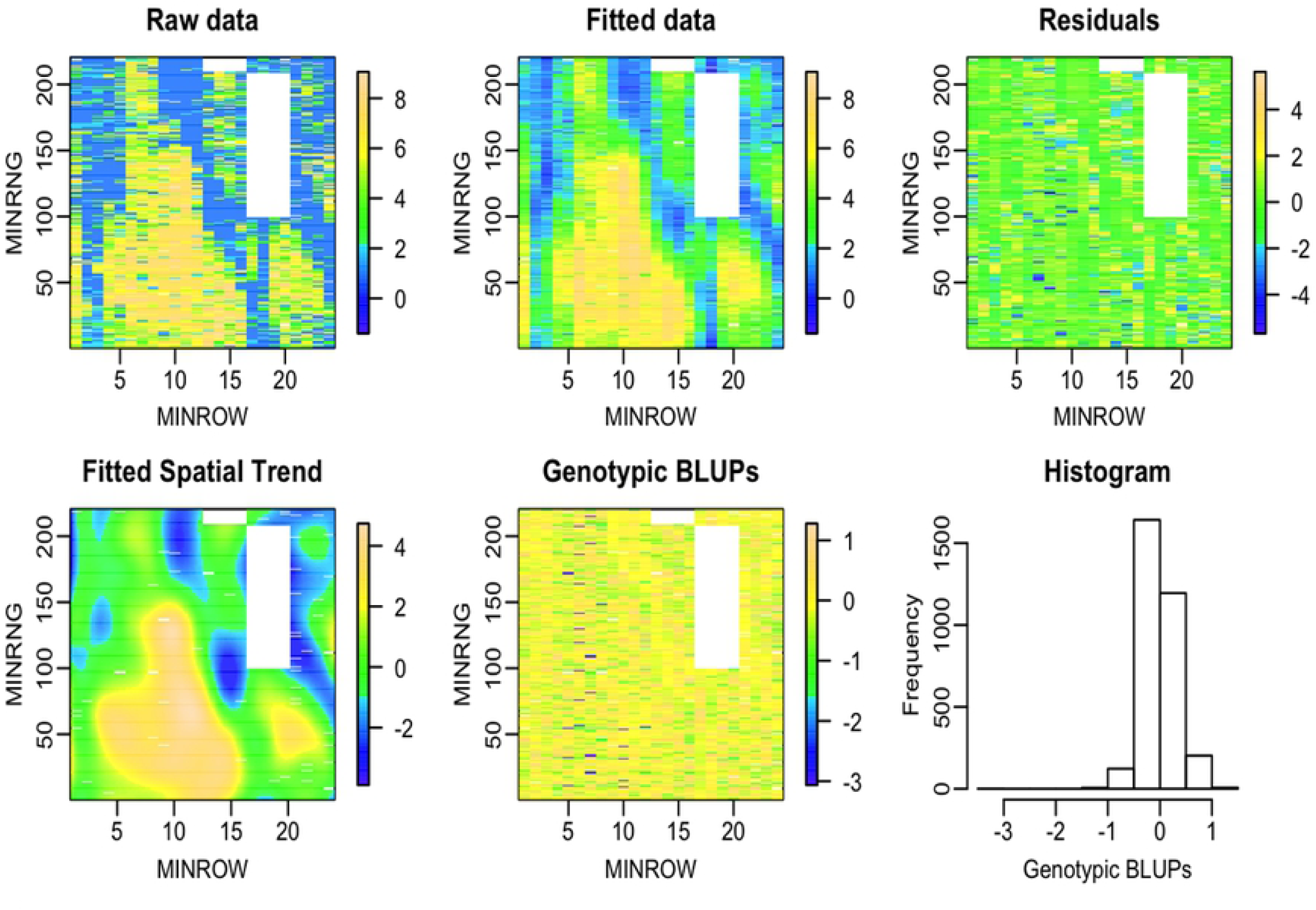
Heatmaps and histogram from data set 3. The rectangle layout is 24 rows (X-axis) by 220 ranges (Y-axis) with a total of 5,280 plots. The white area in the right side of each heatmap are plots without data.

Overall results from the three residual heatmaps obtained from the P-spline model are that spatial patterns or autocorrelations are not fully addressed by the model. These results are different from the two previous reports in which spatial patterns had effectively been removed by 2D P-spline surfaces and residuals for yield in barley [49] and sorghum [50]. We hypothesize that the discrepancy between our results and those previously reported is because yield is a continuous variable and field plot variation was continuous in previously published results. Ordinal IDC scores for small hill plots can be 1 (completely resistant to IDC) in one plot and it’s neighboring plot can be 9 (most susceptible) without a gradual transition between plots. None of the methods developed to date, deal very effectively with these types of data, although the SAR methods provided better RSE values.

Tensor product panelized splines worked very well for hybrid wheat data evaluated in Chilean and Australian wheat field trials [49] as well as sorghum grain yield, and plant height [50]. However, when this method was applied to soybean IDC data sets, two unexpected results were obtained: 1) The effectiveness dimension analysis of the decomposed the model variables reveals that the genotype or line effectiveness accounts for about 90% of the total effectiveness, while the tensor product term “f(ROW):f(RANGE)” accounts for less than 10% of the effectiveness. 2) In contrast to the effectiveness dimension component analysis, the variance component analysis shows that tensor product term “f(ROW):f(RANGE)” accounts for over 90% of the total variance, whereas genotype account for only less than 1%. This suggests a need for development of novel methods with capabilities for analysis of ordinal data obtained from fields that exhibit discontinuous transitions among heterogeneous field plots. We propose that generalized linear, or possibly non-linear, mixed models are most likely to solve these issues.

## Conclusions

The effectiveness of spatial models depends on many factors, such as soil characteristics, weather conditions, field plot operational activities, severity of the spatial variation, and other types of irregular patterns. From the comparison of residual standard error (RSE), R^2^, prediction accuracy, AIC, and observation of their heatmaps generated for eight spatial models, none consistently demonstrate the ability to completely remove the spatial autocorrelation for ordinal data in three data sets. None-the-less, the spatial autoregressive (SAR) approach (with either lag, or error, or Durbin mixed as a covariate) generated residual plots with less evidence of patterns than other models. The tensor product panelized P-splines method applied to simulated data set 1, which had a single circular spatial pattern performed best in this unique situation. As to the computation time and user-friendliness, P-spline implemented int the R software package, SpATS, is the fastest and the easiest to run. Because none of the models could consistently adjust for discontinuous ‘patches’ of ordinal data, there is potential to develop improved novel methods that will be more effective and efficient than are currently available.

## Abbreviations

(IDC): iron deficiency chlorosis
(SAR): geospatial autoregressive regression
(RE): relative efficiency
(OLS w/ RR): ordinary least square with range and row
(AR1): first-order autoregressive

## Supplemental Data Available

Supplemental material is available online for this article. R codes and the data sets used for this research are available online.

## Acknowledgments

This research was supported in part by the Department of Agronomy at Iowa State University, the U.S. Department of Agriculture, Agricultural Research Service, project 5030-21000-062-00D. Mention of trade names or commercial products in this publication is solely for the purpose of providing specific information and does not imply recommendation or endorsement by the Iowa State University or the U.S. Department of Agriculture. Iowa State University and USDA are equal opportunity providers and employers.

## Supporting information

Supplemental Fig 1. IDC field variation heatmap and patterns. (A) Four IDC field variation patterns from heatmaps corresponding to three regions with IDC. (B) Simulated data set 1 with circle pattern. (C) IDC testing field with IDC spatial variability. (D) The three types of the testing locations clustered by soil properties and displayed relative to the first two pricipal components.

